# Efficient fine-tuning of endothelial gene expression by a single Tyrosine to Phenylalanine point mutation in the *VE-cadherin* gene

**DOI:** 10.1101/2024.04.17.589974

**Authors:** Olivia Garnier, Florian Jeanneret, Aude Durand, Arnold Fertin, Sarah Berndt, Gilles Carpentier, Christophe Battail, Donald K. Martin, Isabelle Vilgrain

## Abstract

Cancer and inflammation are associated with vascular diseases that affect endothelial cells (ECs) and alter gene expression. We aimed at understanding whether the site Y^685^ in the cytoplasmic domain of VE-cadherin triggers epigenetic programming *in vivo*. Using our knock-in mice carrying the Y^685^F VE-cadherin mutation, RNA sequencing from lung ECs identified 884 differentially expressed genes (DEG) involved in processes such as cell-cell adhesion, vascular development, and angiogenesis. The 30 DEGs include 22 down-regulated genes (genes encoding cell signalling enzymes, anion transport and lipid metabolism) and 8 up-regulated genes, including the endothelial-specific S1PR1. Analysis of the VEGF/VEGFR2 signaling pathway shows a significant decrease in the expression of pY^1173^VEGFR2 whereas VEGF remains constant, this was consistent with impaired migration, proliferation and protrusive properties of ECs *in vitro*. Co-immunoprecipitation experiments showed that c-Src and Y^685^F-VE-cadherin association which was enhanced in KI compared to WT, resulting in increased in Y^685^F-VE-cadherin phosphorylation at site Y^731^. As a consequence, its partner β-catenin translocates to the nucleus. CHIPS assay showed that FOXF1 binds to the *s1pr1* promoter, leading to increased expression of the S1PR1. In vivo, in the lung vasculature, this process was associated with increased vessel wall thickness and reduced fibrosis. Overall, our findings provide a novel transcriptomic profile triggered by Y^685^F-VE-cadherin ECs for potential insights into therapeutic targets to envisage normalisation of the tumor vasculature.

## Introduction

Cancer and inflammation are associated with vascular disease, which can affect endothelial cells (ECs) and alter gene expression (1). A single monolayer of ECs lines the inside of blood vessels and remains quiescent throughout adulthood. Activation of the endothelium occurs under pathophysiological conditions that lead to either endothelial dysfunction, angiogenesis or changes in the physiology of several organs, particularly those under hormonal control (2,3,4). ECs contact each other via endothelial adherens junctions that express vascular endothelial (VE) cadherin, which is a transmembrane adhesive glycoprotein that is critical for endothelial integrity. Its extracellular domain plays a crucial role in the calcium-dependent adhesion of ECs, while its cytoplasmic tail interacts with the actin cytoskeleton via β-catenin, plakoglobin and p120 to ensure junctional strength (5). The cytoplasmic domain of VE-cadherin regulates endothelial protrusive activity *in vitro*, suggesting that this domain may play a prominent part in invasive processes (6). Considerable evidence has established the important role of tyrosine phosphorylation of the cadherin-catenin complex as a potent mechanism, particularly through the VEGF/VEGFR2 pathway and other inflammatory mediators, all leading to dysregulation of endothelial cell-cell junction stability (7). Importantly, the cytoplasmic domain of VE-cadherin contains nine tyrosine residues that can be potential targets of tyrosine kinases, including one that corresponds to the consensus site (LY^685^AQV) fitting the YxxV/I/L motif for Src family kinases.

In a previous report, we demonstrated *in vivo* VE-cadherin tyrosine phosphorylation and c-Src association in adult mouse tissue under angiogenic conditions (8). This process is dysfunctional in highly angiogenic human glioblastomas and is associated with abnormalities in the tumour capillary endothelial network (9–13). Moreover, in these tissues, as well as under VEGF challenge in HUVECs, the site Y^685^ in the cytoplasmic domain of VE-cadherin has been identified as a target for c-Src. To determine the *in vivo* role of this tyrosine residue, we generated a knock-in transgenic mouse model with a tyrosine-to-phenylalanine (VE-Y^685^F) point mutation in the *VE-cadherin* gene. Consequently, the mice exhibited increased permeability in the female mouse reproductive tract during the hormonal cycle (13,14). A similar increased permeability in the cremaster muscle of male mice was reported by another group of researchers (15). In addition to the VE-cadherin phosphorylation processes associated with endothelial cell-to-cell changes, recent studies have shown that VE-cadherin can significantly contribute to the expression of endothelial-specific genes. This suggests that VE-cadherin can activate signaling pathways leading to transcriptomic programming (16). However, the effect of a phosphorylated form of VE-cadherin on gene expression is not yet known.

Here, we investigated the effect of the Y^685^F-VE-cadherin mutation on the transcriptional activities driven by VE-cadherin. In this manuscript we report our findings that this mutation activates specific transcriptional programs and gene expression profiles in ECs. We identified 884 differentially expressed genes (DEGs) involved in processes such as cell-cell adhesion, vascular development, and angiogenesis. Among these genes, 22 were found to be down-regulated, encoding cell signaling enzymes, anion transport, and lipid metabolism. Those down-regulated genes suggest potential disruptions in crucial cellular pathways. On the other hand, 8 genes were up-regulated, including genes encoding G-coupled transmembrane receptors, such as *s1pr1*. Those up-regulated genes, particularly those encoding G-coupled transmembrane receptors such as S1PR1, indicate the activation of specific signaling cascades. These findings shed light on the intricate molecular mechanisms underlying various biological processes that are important in the pathophysiology of lung diseases. To our knowledge, this is the first report of an efficient fine-tuning of endothelial gene expression by a single Y to Phe mutation in the *VE-cadherin* gene.

## RESULTS

### RNA-Sequencing analysis of lung ECs expressing Y^685^F-VE-cadherin mutant triggers an endothelial-specific-transcription program

To establish a more comprehensive insight into the transcriptional activity of mutant ECs, we performed a bulk RNA sequencing analysis on lung ECs expressing the Y^685^F-VE cadherin mutant and WT. For this purpose, we purified ECs from the lungs of both genotypes at 4-week-old mice. We studied ECs from mice lung to avoid the structural, phenotypic, and functional heterogeneity of ECs depending on the tissue (1). Moreover, we previously published that the murine lung had the highest expression of VE-cadherin compared to other organs (8). The cells were isolated using flow cytometry after collagenase dissociation and single-cell suspensions staining for surface antigens (See Fig.EV1). Lung-specific EC harvests yielded high quality RNA for gene expression analysis. The RNA extracted from the cells was subjected to Next Generation Sequencing analysis (RNA-seq), and the resulting reads were aligned along 94,818 transcripts of the mouse genome (Genomics European Headquarters Bahnhofstr. 804158 Leipzig Germany, Genewiz data. See FigEV 2-6). The number of differentially expressed genes at different fold-change (FC) cut-offs are summarized in Table 1. It is noteworthy that many genes are downregulated in KI compared to WT, suggesting an important function of the mutation in the transcriptomic program.

**Table 1:** SUMMARY OF GENE EXPRESSION CHANGES AFTER Y^685^F MUTATION in VE-CADHERIN.

| \|log2FC \| | 0 | 0.5 | 1.0 | 1.5 | 2.0 | 2.5 | 3.0 | 3.5 | 4.0 |
| --- | --- | --- | --- | --- | --- | --- | --- | --- | --- |
| **Up-regulated** | 8919 | 1946 | 297 | 60 | 15 | 4 | 0 | 0 | 0 |
| **Down-regulated** | 8979 | 4740 | 3039 | 1851 | 851 | 240 | 54 | 19 | 7 |

The global transcriptional changes across the two groups were visualized by a Volcano plot where the log fold change of each gene is represented (x-axis) in function of (-log10) of its adjusted p-value (y-axis). Genes with a corrected p-value less than 0.05 and an absolute log2(FC) greater than 1 are indicated by red dots (Fig. 1A). RNA-seq analysis revealed significant alterations in the gene expression profile, since a total of 884 genes (766 downregulated and 118 genes upregulate in ECs from KI mice) were differentially expressed between the two genotypes. The top 10 significantly enriched pathways (based on p-values) from the gene list are shown in Fig. 1B. The most notably deregulated genes were associated with vascularization, the development of blood vessels and morphogenesis, as well as cell-cell adhesion through plasma membrane adhesion molecules and angiogenesis, including assembly of cell-cell junctions, which are the most deregulated features linked with the endothelial characteristics of Y^685^F-VE-cadherin (Fig. 1AB). More specifically, it is noteworthy that the mutation affected several genes that encode for receptor ligands and transmembrane signaling activities, which include molecular transduction functions, calcium ion binding, and cytokine activities (Fig 1B, C).

**Figure 1.**
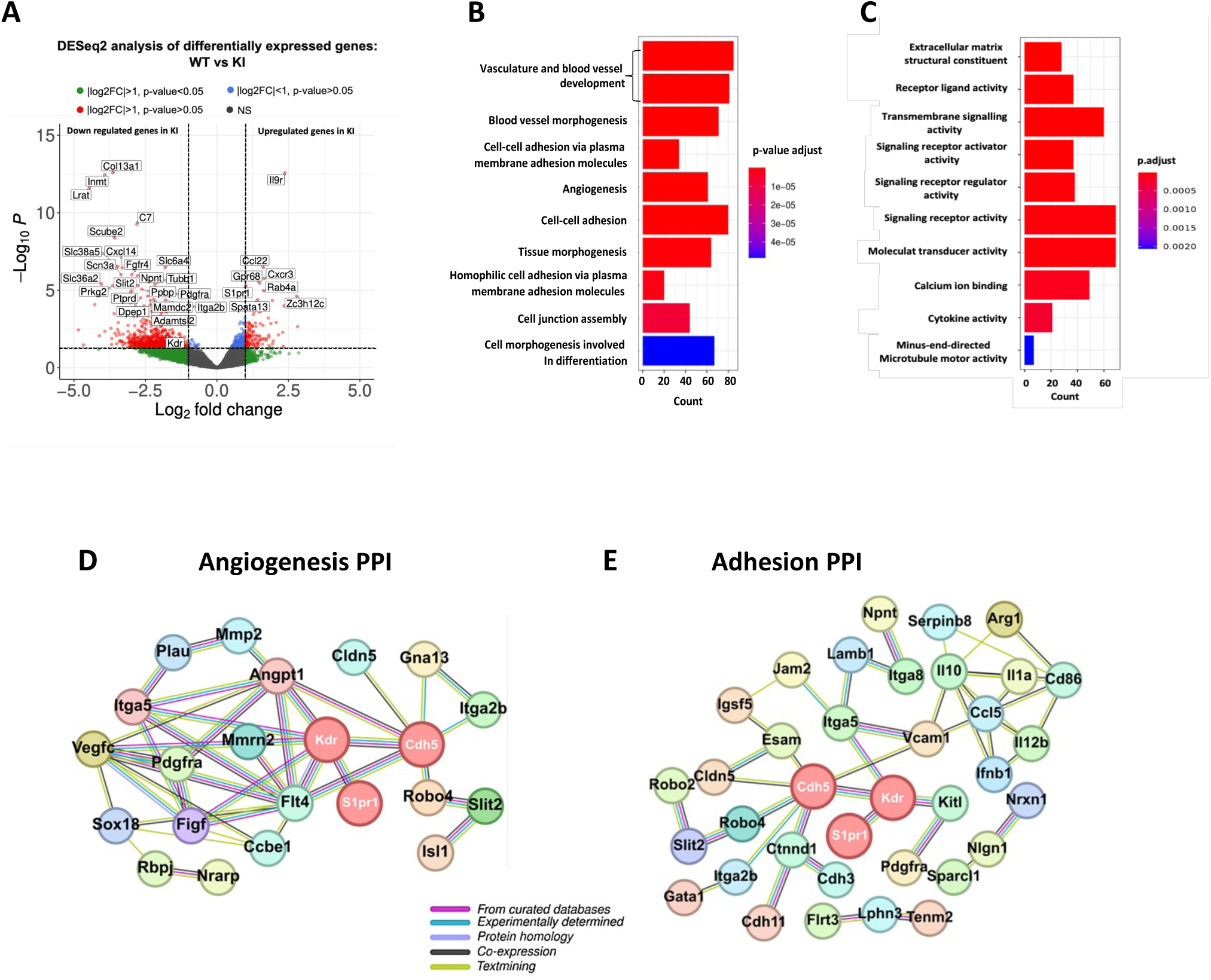
: Transcriptome profile determined by Y^685^F–VE-cadherin mutation demonstrated gene expression involved in angiogenesis and adhesion. **A.** The global transcriptional changes across the WT and the KI mice was visualized by a Volcano plot. Each point represents a gene. The log fold change of each gene is represented on the x-axis and the -log10() of its adjusted p-value on the y-axis. Genes with a corrected p-value less than 0.05 and a absolute log2 (Fold change) greater than 1 are indicated by red dots. **B**. Bar plot shows the top 10 of the Biological Processes (BPs) from the Gene Ontology database enriched by significant genes (corrected p-value < 0.05 and absolute log2(Fold change) > 1). **C.** The DEG involved in cell signalling. **D.** Deregulated protein-protein interaction network of DEGs involved in cell adhesion(BPs). **E**. Deregulated protein-protein interaction network of DEGs involved in angiogenesis (BPs). Protein-protein interaction network was performed using string-db.org software.

To extract biological knowledge, the complete gene expression dataset was submitted to Gene Ontology Enrichment Analysis (GOEA) which revealed the top ten biological processes (BPs) from the enrichment of specific significant genes (corrected p-value < 0.05 and absolute log2(FC) > 1) (Fig. 1D,E). The STRING database was utilized to retrieve interactions among DEGs identified from RNAseq. This network of interacting proteins/genes was then analyzed to assess the coalescence of clusters. The largest protein-protein interaction was found to be associated with the processes of angiogenesis and adhesion. *s1pr1* and *KDR* (VEGFR2) were found to interact each other, and *Cdh5* interacts with *KDR* in cell-cell adhesion and angiogenesis protein-protein networks (Fig. 1D,E). These data suggested the important role of Y^685^F-VE-cadherin in gene expressions that are involved in the main signaling pathway leading ECs to abnormal cell-adhesion and angiogenesis.

### Y^685^F-VE-cadherin inhibits both migration and proliferation with an altered expression of VEGFR2 expression

In addition to providing structural support to ECs, VE-cadherin was known to regulate signaling pathways, including the VEGF/VEGFR2 pathway. Since the transcriptomic analysis indicated that the VE-cadherin mutant had the most significant effect on the angiogenesis pathway, we examined the status of VEGF in the lungs of both WT and Y^685^F-VE-cadherin mouse genotypes. Among the various isoforms of VEGF, VEGF-A165 is the predominant form, of which both VEGF165a and VEGF165b are glycosylated to varying degrees, resulting in molecular weights between 19 and 23 kDa for the monomers and 45 kDa for the dimers. Lung extracts from both genotypes had an equal amount of a 23 kDa protein and a 46 kDa protein as determined by Western blotting which indicated that the expression of VEGF-A was not altered by the Y^685^F-VE-cadherin (Fig. 2A).

**Figure 2.**
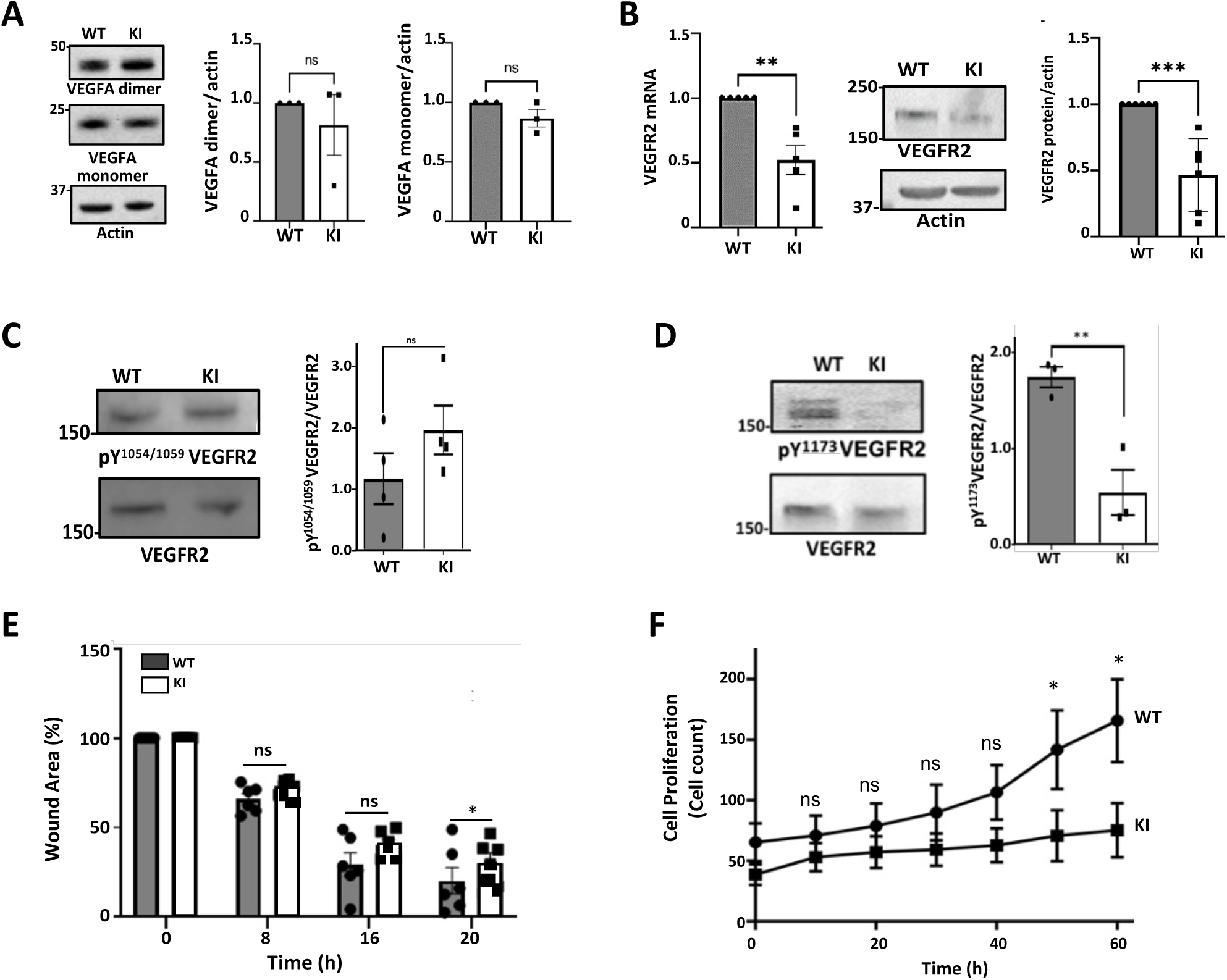
VEGF/VEGF Receptor 2 signaling pathway impair proliferation and migration in mutant EC: **A.** Analysis of VEGFA dimer and monomer by Western blot (n=3 mice per genotype). **B.** VEGFR2 mRNA expression. RT-PCR analysis was performed from isolated EC RNA extracts. VEGFR2 RNA expression level was expressed as the fold-change of WT after normalisation with hprt1. Relative expression of VEGFR2 in ECs from WT and KI mice quantified as the fold-change of WT after normalisation with actin (n=5 mice per genotype). **C**. VEGFR2 autophosphorylation was analyzed using the anti pY^1054/1059^ antibody. The ratio pY^1054/1059^VEGFR2/VEGFR2 was quantified (n=4) **D**. VEGFR2 Y^1173^ phosphorylation was analyzed using the anti pY^1173^ antibody. The ratio was quantified pY^1173^VEGFR2/VEGFR2 was quantified in each genotype (n=4) **E.** Cell culture wound closure assay of ECs from WT and KI. ECs were seeded onto 96-well microplates to reach confluency within 48h. The scratch was made with the Incucyte® 96-Well Woundmaker Tool. The scratch closure was monitored and imaged in 8 h intervals. The percentage of gap area was followed by repeat scanning in the live-cell analysis software until 20h. Magnification (X10 objective) (n=3 mice per genotype). **F.** Proliferation assay for WT and KI cells. ECs were seeded at a density of 1000 cells/mL onto a 96 well plate (n=3 mice per genotype). The proliferation was measured with the Incucyte apparatus for 60h (n=3). Results are expressed as the mean of triplicate determinations and the experiment was repeated three times with similar results. All immunoblots were carried with 30 μg of total proteins and protein level was normalized with actin. Molecular mass standards (in kDa) are shown at left. The relative expression of the protein were measured by densitometry of autoradiographs using ImageJ software (NIH, Bethesda, MD). Statistical analysis was performed using Student’s t-test or Mann-Whitney test as described in Methods, *p-value<0.05, ** p-value<0.01, **** p-value<0.0001. Non significant ns)

Since VEGFR2 is expressed almost exclusively in ECs and because our transcriptomic data shown that the gene expression of VEGFR2 was -1.773 with an adjusted p-value of 0.0084, we attempted to detect VEGFR2 mRNA and protein levels in ECs from both genotypes. The results showed that the relative expression of VEGFR2 mRNA was decreased by 50% in the KI mouse compared to the WT (p-value= 0.012) (Fig.2B). When normalized to actin, the VEGFR2 protein level was also significantly decreased in KI cells (p-value=0.02) (Fig. 2C). Since VEGFA was expressed at the same level in WT and KI mice and is an endogenous ligand for VEGFR2, it was expected that VEGFR2 would be permanently activated in lung ECs. There are two important tyrosine sites, Y^1054^ and Y^1059^, which are required for full activity of the receptor tyrosine kinase. Indeed, previous data have shown that mutation of Y^1054^ and Y^1059^ to phenylalanine residues results in loss of kinase activity (17). Therefore, we measured the degree of activation of VEGFR2 using an antibody directed against pY^1054/1059^ VEGFR2. Our results showed a trend towards higher levels in the KI, but not significantly different from the WT, suggesting that the VEGFR2 has the same kinase activity in both genotypes. This result is in line with previous literature which has shown that phosphorylated Y^1054/1059^ are required for the maximal VEGFR2 activity but not essential for the VEGF-A-induced proliferation in ECs (18). Another important site for VEGFR2 is the tyrosine Y^1173^, which is an essential binding site for PLCγ via the SH2 domain when it is phosphorylated. This process is involved in VEGF-A-induced cell migration and proliferation (19). Using an anti-Y^1173^ VEGFR2 antibody, we demonstrate that the phosphorylation of the site Y^1173^ was reduced by 30% in KI mice (0.53 ± 0.23) compared with WT (1.71 ± 0.10) (p=0.009) (Figure 2D). This prompted us to determine the migration and the proliferation properties of isolated ECs from KI and WT lungs. First we analyzed the migratory activity of the ECs using the wound closure assay which is one of the basic readout in angiogenesis (20). The wound closure assay was performed using the Incucyte® Cell Migration Kit (Sartorius) and imaged over 8 hour time period until 20 h (Fig.2E). The results indicated that within 20 hours, WT and KI ECs gradually filled the available space in a time-dependent manner starting at 8 hours, but for KI ECs to migrate slower than WT ECs so that after 20 hours there was a significant difference in wound area between the KI and WT ECs (WT 13.24% ± 5.17; KI 22.45% ± 5.20, p-value=0.034).These findings are consistent with our data that showed inhibition of VEGFR2 expression in the Y^685^F-VE-cadherin mutant. Moreover, our results confirm the previous suggestion that the degree of phosphorylation of Y^1173^VEGFR2 plays a role in the collective migration of ECs, angiogenesis and vascular modelling (21).

We next examined the proliferation rate of ECs from both genotypes. The proliferation assay and cell proliferation were performed by the Incucyte Basic Analyzer software. The results in Fig2F show that ECs with mutated VE-cadherin did not initiate cell division during the tested time period. In contrast, ECs from WT mice begin to proliferate significantly within 40-60 hours, which corresponds to the slow turnover rate observed in ECs from other organs (WT 141.85 ± 32.52 vs KI 70.66 ± 14.86, p=0.040 for 50h; WT 165.71 ± 34.14 vs KI 75.22 ± 24.25, p=0.037 for 60h) (Fig. 2F). Altogether, our results strongly suggest that the mutation of VE-cadherin had an impact on VEGFR2 autophosphorylation site Y^1173^ and, as a consequence, a significant reduction in both the proliferation and migration of ECs.

### Y^685^F-VE-cadherin ECs exhibit reduced angiogenic properties in the 3D Fibrin Bead Assay

Angiogenic properties of ECs from both genotypes were determined using the fibrin bead assay, a classic test of angiogenesis that mimics sprouting in a complex 3D model incorporating a matrix that resemble the native *in vivo* environment (20). ECs were first allowed to adhere to collagen I-coated beads to generate a monolayer that mimicked the vessel wall, then these EC-coated beads were embedded into a fibrin gel with human lung stromal cells (e.g. fibroblasts). After a few days in culture, the elongation of pseudo-vessels appeared as tree structures. The images of this original cellular organization were acquired through phase contrast microscopy (Fig. 3A), and the tree structures had been broken down as Junctions and Extremities as well as Segments, Branches and Anchorage Junctions and meshes to quantify those morphological parameters (Fig. 3B). Image analysis was performed using a program developed for the ImageJ software (22). A representative map of the detected object is depicted in Fig 3C with the legend in Fig. 3D. Table 2 summarizes the complete analysis of angiogenic properties from 88 beads containing WT ECs compared with 51 beads containing KI ECs after 4 days of culture (Fig. 3E,F).

**Figure 3:**
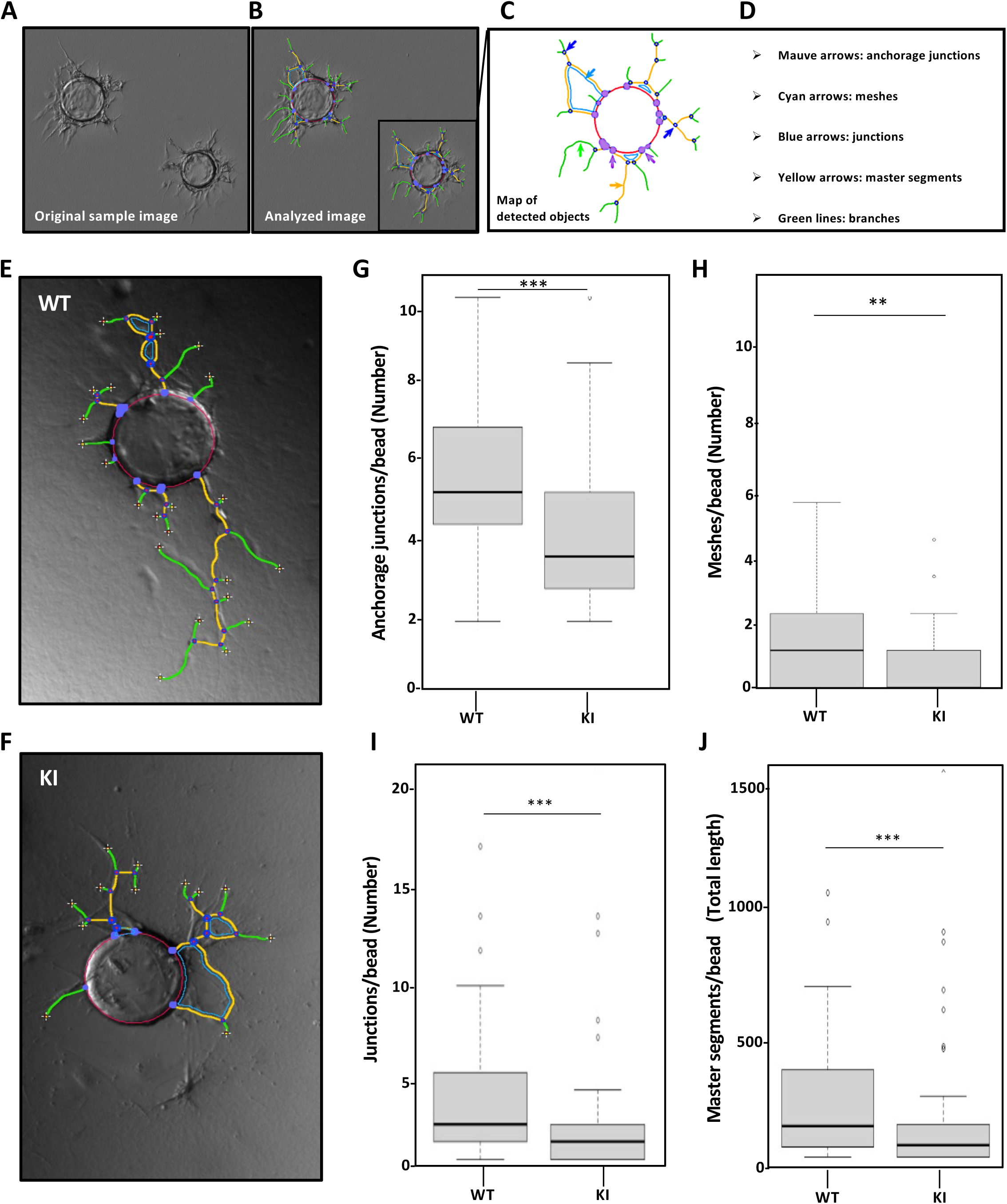
Y^685^F-VE-cadherin EC exhibit reduced angiogenic properties in the 3D Fibrin Bead Assay. **A-D.** Vectorial objects characterized and quantified by the software analysis for the WT and KI cellular trees. **E-F**. Representative images of WT and KI EC coated bead after 4 days in fibrin. **G-J.** Quantification of morphometrical parameters of the capillary network was performed by a computerized method on images taken on day 4. Representative parameters measured were: number of anchorage junctions, number of meshes, number of junctions, and number of master segments per bead. 88 beads for WT and 51 for KI were quantified. All bar graphs showed mean ± SEM. Statistical analysis was performed using Mann-Whitney test as described in Methods, *p-value<0.05, **p-value<0.01, ***p-value<0.0001. Quantification of morphometrical parameters of the capillary network was performed using a program developed for the ImageJ software.

**Table 2:** Vectorial objects characterized and quantified by the software analysis for the WT and KI ECs.

| Vectorial element | Short definition | WT (mean) ± SEM | KI (mean) ± SEM | p-value |
| --- | --- | --- | --- | --- |
| *Anchorage Junction* | Junction linking a *Branch* or a *Master Segment* to a *Sphere* limit | 5.46 ± 0.25 | 3.9 ± 0.30 | 0.0001 |
| *Junction* | Group of *Nodes* forming a bifurcation | 3.18 ± 0.35 | 2.19 ± 0.58 | 0.0009 |
| *Total Master Segment length* | Sum of length of binary lines remaining after removal of branches | 234.96 ± 26.31 | 167.33 ± 46.12 | 0.0007 |
| *Meshes* | Area enclosed by *Master Segments*, or *Master Segments* and *Sphere* limits | 1.045 ± 0.119 | 0.647 ± 0.15 | 0.01 |

Anchorage junctions are the number of junctions between the master segment and the spherical border. Consistent with the wound assay described above, it is noticeable that these are significantly reduced in KI ECs, indicating that these cells have less protrusive activity compared to WT ECs. (Fig. 3G). In addition, and consistent with this, the EC in 3D gel grow according to a meshed network that was quantified and showed that this level of cellular organization was also decreased in KI ECS when compared to WT (Fig.3H). All of the ECs spontaneously aligned and branched, but the junctions forming a bifurcation appeared in a relatively lower quantity in KI ECs than in WT (Fig. 3I). Importantly, the ECs from KI exhibited a significantly reduced length of master segments, which indicate that these cells do not migrate like ECs from the WT (Fig. 3J). Altogether, these results revealed that all the parameters measured using this fibrin bead assay were dramatically reduced for KI ECs as compared to the WT ECs. Therefore, based on these experiments, it can be concluded that the VE-cadherin point mutation results in a strong impairment of the protrusive activities of the protein as well as the migratory parameters of ECs.

### Y^685^F-VE-cadherin is associated with active c-Src leading to adhesion properties and EC morphometric shape changes

c-Src is basally inactive in most normal tissue due to autoinhibition by intramolecular interactions between its c-Src homology 2 domain (SH2) and a phosphotyrosine (Y^527^) in the C-terminal region. Phosphorylation of c-Src at Y^527^ is catalyzed by the C-terminal Src kinase (CSK), a cytoplasmic tyrosine kinase that has this high substrate specificity which leads to c-Src inactivation (23). Moreover, it was previously described that in non-migrating cells, CSK was found to co-localize with c-Src supporting that Src inactivation mediated by CSK plays an important role in the regulation of cell adhesion (24). Since c-Src function was reported to be attenuated by CSK overexpression in murine colon cancer cells (25), we examined the relative status of both tyrosine kinase expression in the lungs of WT and KI mice. We performed an immunoprecipitation of CSK from lungs and probed with the antibody against c-Src (Figure 4A). Interestingly, we found that the interaction of CSK with c-Src was higher in WT mice than in KI, suggesting that c-Src activity was greater in KI than in WT. To investigate c-Src activity in ECs from lungs, we used the anti-phosphoY^418^c-Src to monitor active kinase since phosphorylation at site Y^418^ is required for full activation of c-Src. The results showed that the level of pY^418^c-Src was higher in KI (1.29 ± 0.09) than in WT (0.77 ± 0.07) (p-value= 0.02) (Fig. 4B). Therefore, we conclude that c-Src activation was dependent on CSK expression and was more active in KI than in WT.

**Figure 4:**
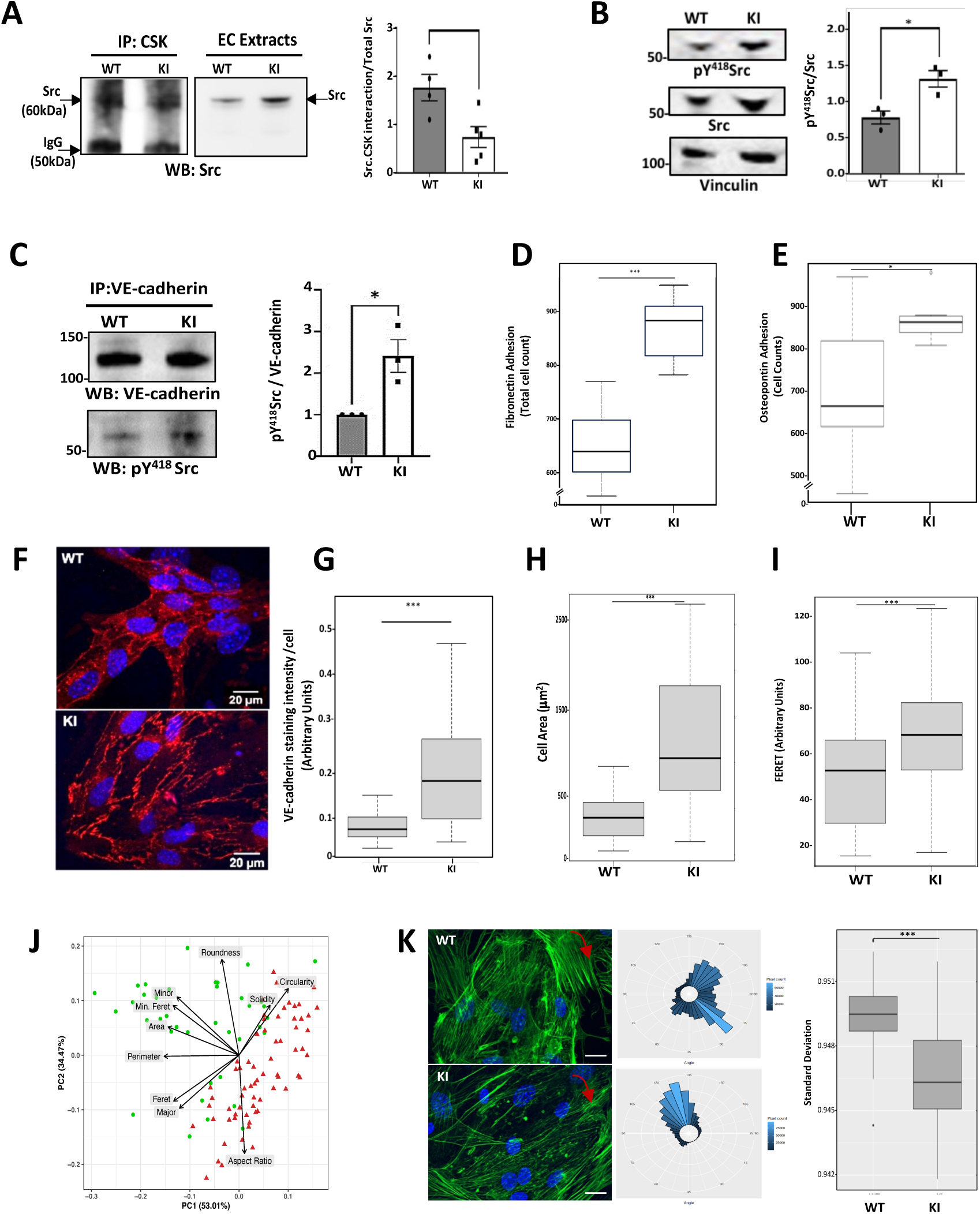
Active Src is a partner of Y^685^F-VE-cadherin leading to adhesion properties and EC morphological changes. **A.** Representative CSK immunoprecipitate from lung mice extract (100 μg) probed with the anti Src antibody (n=4 for WT and n=5 for KI). **B.** Western blot analysis of pY^418^ Src and Src were performed using 5 μg of protein extracts from the isolated ECs from WT and KI mice. Vinculin was used as loading control (n=3 mice per genotype). Molecular mass standards (in kDa) are shown at left. On the right is illustrated the relative expression of the pY^418^Src/Src measured by densitometry of autoradiographs using ImageJ software. **C.** Representative VE-cadherin immunoprecipitate from lung mice extract (100 μg) probed with the anti phosphoY^418^Src antibody (n=3 mice per genotype). On the right handside is illustrated the relative expression of the pY^418^Src/VE-cadherin measured by densitometry of autoradiographs using ImageJ software. **D.** Adhesion onto FN-coated coverslip: Cells were seeded at 1500 cells/96 wells onto fibronectin coated plate (10 μg/mL) and incubated for 30 min and stained with Hoescht. The nuclei were counted for each well (n=6). All images were quantified using ImageJ software. **E.** Adhesion onto Osteopontin-coated plates: Cells were seeded at 1500 cells/96 wells onto OPN-coated plate (0.3 μg/mL) and incubated for 30 min and stained with Hoescht. The nuclei were counted for each well (n=6). All images were quantified using ImageJ software. **F.** Representative immunostaining of VE-cadherin on isolated ECs. Hoescht was used to stain cell nuclei. Magnification (X63 objective). Scale bars: 20 μm. **G**. Quantification of VE-cadherin staining at the membrane. **H.** Cell Area quantification from WT and KI. Images are representative of 3 mice per genotype (WT n=29; KI n=17). **I. J.** Cell area was quantified**. K.** Orientation of the cell actin filaments under flux (Left panel). Representative images of EC from 3 mice per genotype (WT n=25; KI n=47) Hoescht was used to stain cell nuclei. Magnification (X63 objective). Scale bars: 20 μm. Representative circular plot of actin filament (Center). Comparison of standard deviation between WT and KI (Kruskall.Wallis test, p-value = 2.342 10^-6^) (right panel). All bar graphs showed mean ± SEM. Statistical analysis was performed using Mann-Whitney test as described in Methods, otherwise as stated in the text (*p-value<0.05, **p-value<0.01, *** p-value<0.0001) .

As we reported previously, there is a constitutive association between VE-cadherin and c-Src in mouse organs in vivo, including the ovary and uterus (8). Therefore, we investigated whether an association between pY^418^c-Src and VE-cadherin was detected in both genotypes *in vivo*. To do so, we performed immunoprecipitation on lung lysates using a VE-cadherin antibody, followed by immunoblotting with an anti-pY^418^c-Src antibody. The immunoprecipitate analysis revealed the presence of pY^418^c-Src in both KI and WT mice. However, the level of pY^418^c-Src was significantly higher in KI mice (0.60 ± 0.035; p = 0.0269) compared to WT mice (0.35 ± 0.0643) (Fig 4C). These results provide strong evidence that c-Src remains localized at the EC junctions. Moreover, the Y^685^F-VE-cadherin mutation enhances the interaction between pY^418^c-Src and VE-cadherin.

Because c-Src plays a crucial role in regulating cell-extracellular matrix (ECM) proteins adhesions, we next investigated the initial adhesion of ECs to fibronectin (FN), a classical adhesion-promoting protein for ECs (26) and osteopontin (OPN) another extracellular matrix protein important for angiogenesis (27). We seeded suspensions of ECs isolated from the lungs of both genotypes (1500 cells/mL) onto coverslips coated with FN (10μg/mL) and with OPN (3μg/mL) and counted the nuclei of the attached cells after 30 minutes. A significant difference in EC adhesion was observed in KI ECs for both substrates indicating a stronger interaction when the active c-Src associated with VE-cadherin was at highest level (p-value=0.00028) (Fig. 4D,E). Collectively, these data confirmed that ECs from KI mice adhere more rapidly than those from WT mice at the first cell-substrate contact (28). Furthermore, this is also consistent with the reported literature showing that c-Src in normal cells is concentrated at sites of membrane-substrate interaction (29). Another crucial role attributed to the kinase c-Src is the regulation of endothelial cell-cell junctions. To gain a better understanding of how the association between c-Src and VE-cadherin affects the quality of endothelial junctions, we conducted a comprehensive immunofluorescence analysis of VE-cadherin. In ECs from WT, the staining pattern consisted of a thin and continuous line highlighting the margins of each cell. These findings are similar to previously reported VE-cadherin localization to filopodia-like structures connecting two adjacent ECs. In contrast, in the mutated ECs, the fluorescence signal along cell margins was more discontinuous and in many areas more diffuse than in the WT with a distribution in multiple punctate accumulations (Fig 4F). Quantification of fluorescence intensity at the membrane revealed that the mutant ECs had a significantly higher amount of VE-cadherin at the membrane than the WT (Figure 4G). We then analyzed the morphometric parameters of the ECs, including cell area, which was greater in KI ECs compared to WT ECs (Fig. 4H, see also Fig EV9). In addition, there was an increase in the magnitude of the Feret’s diameter (Fig.4I). All the significant parameters characterizing the ECs in the principal component analysis (PCA), including perimeter vector, area, minor axis and major axis are shown in Fig 4J. The graph presented illustrates two distinct clusters of data points, each representing the KI/WT group. These clusters indicate a noticeable difference between the two groups. These PCA results allow the inference that the Y^685^F-VE cadherin-based endothelial junctions exhibit a distinctive cellular morphology characterized by elongated cells with increased adhesive properties, both towards the ECM and to neighboring cells.

*In vivo,* blood pressure measurements were altered in the Y^685^F-VE cadherin mice, demonstrating endothelial dysfunction (See Fig. EV 10). We therefore wondered whether in vitro ECs of both genotypes would be affected by experimental changes in blood flow. Since VE-cadherin is linked to the actin cytoskeleton, we followed the actin network under shear stress to see if adaptive adjustments occurred (30). To achieve this, we used a model that applies uniaxial flow to cells at the periphery, since capillaries in the lung are typically exposed to minimal shear stress compared to larger arteries. The ECs were seeded onto coverslips in a 24-well plates and subjected to shear forces using a rotary shaker for 72 hours at a rotation frequency of 110 rpm, corresponding to shear forces of 3.3 dynes/cm2 (31). Actin filament orientation was detected using phalloidin-FITC. Figure 4K (middle panel) shows the analysis of the structural variations measured from WT ECs (n=27) compared with KI ECs (n=42), and representative images of the ECs (left panel). The position of the actin filaments is quantified in the polar bar plot (Fig. 4K, middle panel), which shows a significantly lower angular dispersion of actin in KI cells. This is further quantified as a smaller standard deviation, as shown in the box plot of the standard deviation (p-value = 2.342 x 10^-6^, Kruskal-Wallis test) (Fig. 4K, right-hand side). These results confirm that KI ECs undergo structural changes compared to WT ECs and support the hypothesis that the Y^685^F-VE-cadherin mutation enhances the ability of the actin cytoskeleton to associate more with the ECM, which is consistent with our results that lung ECs from KI mice adhere more rapidly than those from WT mice, and published information of the increased adhesive properties of ECs (32).

### Y^685^F-VE-cadherin increased VE-cadherin phosphorylation at site Y^731^ leading to β-catenin translocation to the nucleus

Considering our result that Y^685^F-VE-cadherin is associated with active c-Src, we hypothezised that this association may result in the tyrosine phosphorylation of VE-cadherin at additional sites. One site of interest is the amino acid residue Y731, which has been identified as a binding site for β-catenin (34). To investigate this site, we used immunoblotting with the anti-pY^731^-VE-cadherin antibody, which showed that the Y^731^site was phosphorylated in both genotypes, but the phosphorylation was markedly higher in KI (1.48 ± 0.14) than in WT (0.82 ± 0.13) (p-value=0.041) (Figure 5A). Consequently the phosphorylation of this site would, at least in part, reduce the binding of β-catenin to VE-cadherin. Therefore, we investigated the association between VE-cadherin and β-catenin in both genotypes by immunoprecipitating VE-cadherin using an anti-VE-cadherin antibody. As anticipated, we observed a significant decrease in the ratio of VE-cadherin to β-catenin in KI mice (0.798 ± 0.098) compared to WT mice (1.136 ± 0.092) (p-value=0.0369) (Fig. 5B). In line with the Cheresh laboratory’s findings, our results confirmed that β-catenin molecules were removed from the pY^731^VE-cadherin adherens junction complex. Since β-catenin plays a crucial role in regulating gene expression within the nucleus (35), we examined whether the amount of β-catenin in the nuclei of lung ECs was modified depending on the genotypes. To achieve this, we analyzed the nuclear fraction purified from KI and WT mice by immunoblotting with an antibody to β-catenin (90 kDa) and with an antibody directed against histone H3 (15-17 kDa), which is a marker of nuclei and was also provided a loading control. The results show a significant increase in the β-catenin/H3 ratio in the nuclear fraction of the KI mice (1.11 ± 0.14) compared to the WT mice (0.69 ± 0.10) (p-value=0.040) (n=7 mice per genotype) (Figure 5C). Collectively, these results demonstrate that c-Src induces phosphorylation of VE-cadherin at site 731, which partially impairs the binding of β-catenin to the adherens complex, an effect that is more pronounced in KI mice.

**Figure 5:**
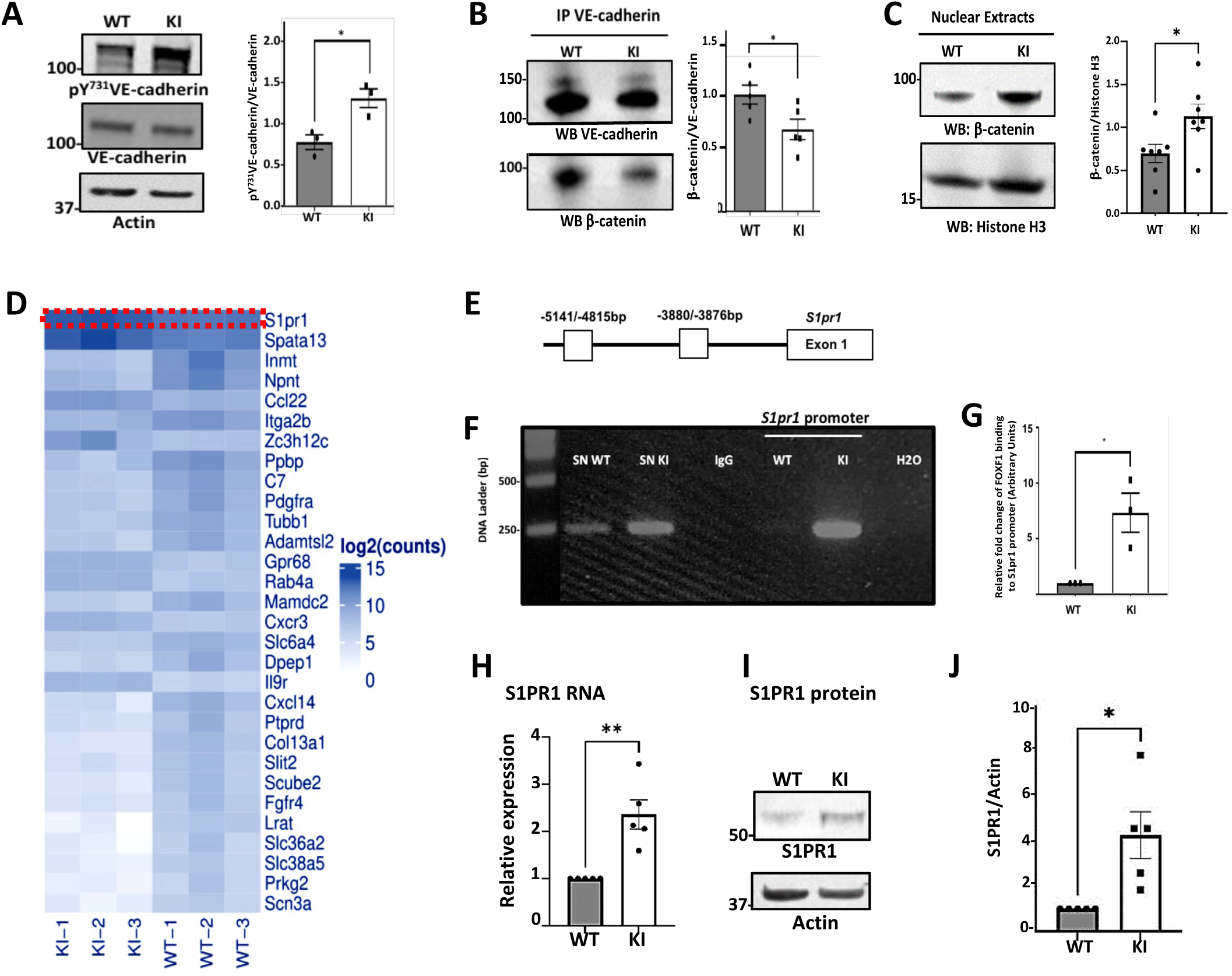
Increased VE-cadherin phosphorylation at site Y^731^ leads to β-catenin translocation to the nucleus, which activates FOXF1 that binds directly to the *s1pr1* promoter and increases S1PR1 levels. **A.** Analysis of pY^731^VE-cadherin and VE-cadherin in WT and KI using 30 μg of lung extracts. Actin was used as loading control (n=3 mice per genotype). On the right is illustrated the relative expression of the pY^731^VE-cadherin/VE-cadherin measured by densitometry of autoradiographs using Image J software. **B.** Representative western blot of VE-cadherin immunoprecipitate from lung mice extract (100 μg) probed with β-catenin antibody (n=5 mice per genotype). On the right handside is illustrated the relative expression of the β-catenin /VE-cadherin measured by densitometry of autoradiographs using Image-J software. **C**. Analysis of nuclear fraction from WT and KI mice with β-catenin antibody using 16µg of proteins. Histone H3 was used as loading control (n=7 mice per genotype). Molecular mass standards (in kDa) are shown at left. On the right is illustrated the relative expression of the β-catenin /Histone H3 measured by densitometry of autoradiographs using Image J software. Statistical analysis was performed using Mann-Whitney test (***p-value<0.001) as described in Methods. **D.** Heatmap of the top 30 differentially expressed genes ranked according to their average expression value across all samples. The differentially expressed gene *s1pr1* is highlighted by the red dotted line. **E**. Schematic drawing of the mouse *s1pr1* promoter region with potential FOXF1 DNA binding sites (white boxes). FOXF1 protein binds to the −5141/−4815-bp and −3880/−3876-bp of s1pr1 promoter regions**. F.** FOXF1 cross-linked protein/DNA complexes were immunoprecipitated with rabbit anti-FOXF1. The same quantity of DNA was loaded onto the agarose gel for each fraction (Supernatant (SN) and s1pr1 promoter regions bound to FOXF1). RT-qPCR was performed with primers designed to capture the FOXF1-binding site near the transcription start site of the *s1pr1* promoter, negatively and positively controlled by IgG and total input without immunoprecipitation. **G**. The relative fold change of the binding of FOXF1 on s1pr1 promoter regions between WT and KI (n=3 mice per genotype) was quantified using Image J software. Molecular mass standards (in kDa) are shown at left. **H.** Relative expression of S1PR1 mRNA in ECs from WT and KI mice quantified after normalisation with hprt1. **I.** Immunoblotting analysis of lung protein extracts (30 mg) was performed using the S1PR1 antibody and normalized with actin (n=5 mice per genotype). **J.** Relative expression of S1PR1 /actin from WT and KI mice measured by densitometry of autoradiographs using ImageJ software. Statistical analysis was performed using Mann-Whitney test (*p-value<0.05, **p-value<0.01, ***p-value<0.0001).

### FOXF1 which binds directly to the S1PR1 promoter increasing S1PR1 mRNA and protein levels

Our results that demonstrated a partial translocation of β-catenin to KI nuclei supports an hypothesis that this translocation is likely to be responsible for the differential gene expression observed in our transcriptomic analysis of ECs from KI and WT (35). In support of this hypothesis, Figure 5D shows the heat map of the top 30 genes sorted by their adjusted p-value and their log2-transformed expression values across samples. The reproducibility among biological triplicates demonstrates the fidelity of the identification, isolation, and profiling of lung-specific ECs in both WT and KI mice. The first gene in the list is *s1pr1*, which is highlighted due to it having the highest adjusted p-value (p>10-12). This gene encodes sphingosine-1-phosphate receptor 1, a G-coupled transmembrane receptor that is highly expressed in ECs (36). Previous literature has shown that endothelial-specific S1PR1 knockout mice exhibit vascular leakage in several organs, including the lung. (37). Additionally, previous reports have identified the transcription factor FOXF1, which is highly expressed in resting adult lung ECs, as a potential transcriptional regulator of the *s1pr1* gene (38–40). Since FOXF1 has been shown to maintain endothelial barrier function, we hypothesized that FOXF1 could directly regulate the *s1pr1* promoter in both mouse genotypes to modulate endothelial junctions and VE-cadherin. Previous reports in the literature have shown that FOXF1 binds specifically to the -3880/-3876 base pair (bp) and -5141/-4815 bp regions of the *s1pr1* promoter. (39) (Fig. 5E). Thus, we used a chromatin immunoprecipitation assay (CHIP) from mouse lung nuclei extracts from KI and WT to determine whether FOXF1 protein could directly bind to *s1pr1* promoter. We chose the −5141/−4815-bp DNA fragment to perform RT-qPCR with primers (see table Fig.Extended 1) designed to capture the FOXF1-binding site near the transcription start site of the *s1pr1* promoter. EconoTaq Plus Green was used according to the following cycles: 3 minutes at 95°C, 34 cycles: 95°C 15 seconds; 62°C 15 seconds; 72°C 20 seconds, 72°C 10 minutes. The DNA concentration was measured in order to allow us to load the same amount of DNA (400ng) onto the agarose gel for analysis. The result clearly showed that the FOXF1 protein binds directly to the *s1pr1* promoter, with a significant seven-fold increase in KI ECs compared to WT ECs (fold-change 7 ± 1.76, p-value=0.023) (Fig.5 F,G). The binding specificity was confirmed by immunoblotting FOXF1 in the CHIP samples (data not shown). These results confirm that FOXF1 directly regulates *s1pr1* gene transcription in KI mouse ECs. To expand this data, we performed a real-time PCR analysis of S1PR1 mRNA in both genotypes. The results showed a significant increase in expression in KI, with an average fold change of 3.26 ± 0.310 (p-value=0.011) (Fig. 5H). In addition, using immunoblotting and normalization to β-actin, we found that S1PR1 protein was significantly upregulated in KI mice compared to WT mice (p-value=0.034) (Figure 5I,J). In conclusion, the transcription factor FOXF1 binds to the *s1pr1* promoter and activates *s1pr1* gene transcription in KI mice. This is supported by an increase in both mRNA and protein, indicating a potential regulatory mechanism involving FOXF1 and S1PR1 in the context of VE-cadherin mutation.

### Lungs from Y^685^F-VE cadherin KI mice have reduced number of arteries with thicker walls

Since S1PR1 is highly expressed in quiescent adult lung ECs and has been shown to play an important role in the maintenance of endothelial barrier function and vascular maturation (36), we investigated whether differences in lung morphology exist, focusing on the vascular pattern of the two genotypes. Because S1PR1 expression was restricted to the alveolar capillaries and arterioles, we counted the small pulmonary arteries with a comparable diameter of 75-150 μm. The data revealed a significant decrease in the number of vessels in the KI group (287 vessels) compared to the WT group (411 vessels) (p-value=0.036) (4 mice per genotype)(Fig. 6A,B). The data revealed a significant increase in the morphometric standard medial thickness of pulmonary arteries in KI mice compared to WT mice (41.39 ± 6.48 µm vs. 26.87 ± 1.08 µm, p-value=0.049) (Fig. 6C). In addition, the ratio of medial area to luminal area was higher in KI mice than in WT mice (2.99 ± 0.506 vs. 1.49 ± 0.065, p-value=0.000013), supporting the above results (Fig. 6D). This finding prompted further investigation into whether there was a change in permeability between KI and WT. Collagen is a structural protein present in connective tissue and is known to accumulate in fibrotic lungs, which is associated with an increase in permeability. We performed an histological analysis of collagen in the lungs of WT and KI mice, with the results showing less collagen accumulation around the walls of small pulmonary arteries in the lungs of KI mice compared to WT mice (KI mean= 53206 ± 338; WT mean =70039.02 ± 3142; p-value=0.0008) (Fig.6E,F). The data are consistent with our transcriptome analysis, which showed a significant decrease in the expression of Col XIII, which is involved in cell-matrix and cell-cell adhesion, in KI mice (Log2 FC -3.646; p=value = 2x10^-13^). Therefore, the correlation between S1PR1 expression and the increase in vessel wall thickness, as well as the reduction in collagen, suggests that the KI mice may be less sensitive to fibrosis due to the strengthening of endothelial barrier integrity by S1PR1. Our measurement of the wet-to-dry lung weight ratio indicated the absence of pulmonary edema, further supported these findings (Fig.EV8).

**Figure 6.**
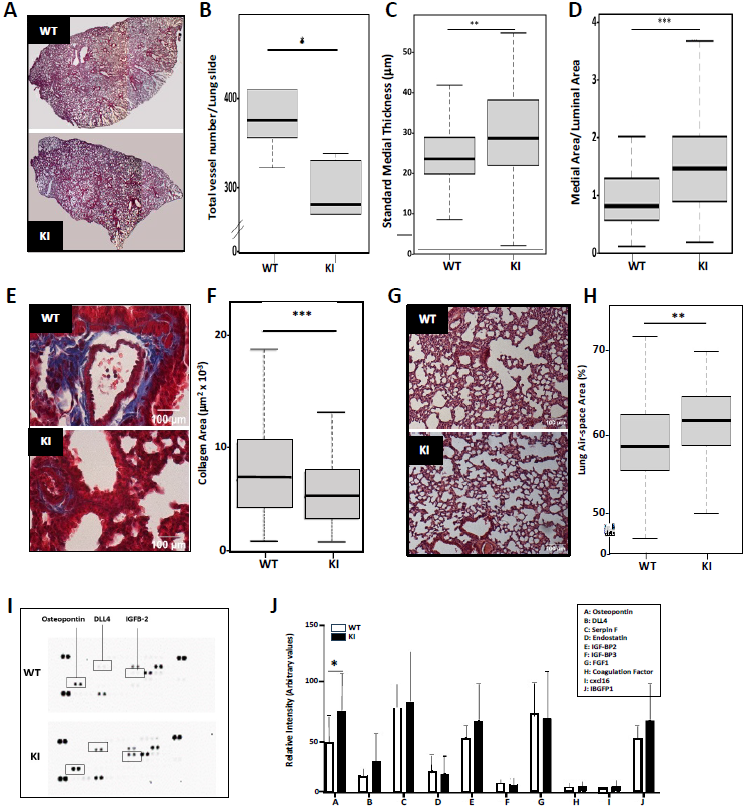
The Y^685^F-VE cadherin mutation results in abnormal lung features, as well as changes in the content of BAL fluid. **A.** Representative H&E images of lung sections from KI mouse vs WT mouse used to count the vessels. Magnification (x40 objective), scale bar 100µm. Images are representative of 4 mice per genotype. **B**. The small pulmonary arteries of comparable diameter (75-150 μm) were counted and showed a decrease in KI mice. Images are representative of 4 mice per genotype. **C**. Morphometric analysis of the arterial standard thickness. **D**. Morphometric analysis of medial/luminal area per vessel. **E**. Representative hematoxylin-eosin images of lung sections showing collagen disappearence in KI mice. Magnification (x10), n=3 lung for each genotype. Scale bars: 100 μm. **F.** Collagen was quantified using ImageJ software. **G**. Representative H&E lung sections displaying alveolar air spaces. Magnification (x10 objective), n=3 lung for each genotype. Scale bars: 100 μm. **H.** Alveolar air-space area quantifications are expressed as % (WT lung slices n=56, KI n=55) (p-value=0.0059). **I.** Protein array was performed with « Mouse Angionesis Array » in the BAL fluid according to the manufacturer’s instructions. The same amount of proteins was applied onto the membranes. **J**.The relative intensity of the array spots between the two genotypes were quantified using ImageJ software (n=4 mice per genotype). All bar graphs showed mean ± SEM. Statistical analysis was performed using Mann-Whitney test as described in Methods, otherwise as stated using Wilcoson test (*p-value<0.05, **p-value<0.01, ***p-value<0.001, ****p-value<0.0001).

Next, because the lungs play a crucial role in gas exchange due to their high vascular density, we investigated whether there were any differences in lung morphology between WT and KI mice. To do this, we determined the air space in lung sections from both genotypes using haematoxylin-eosin images. Light microscopic examination suggested that the KI mice had an enlarged airspace, which was confirmed by quantitative histomorphology. A significant increase in the alveolar size of KI mice (KI 61.93 ± 0.94%, n=55) was observed vs WT (58.71 ± 0.70%, n=56) (p-value=0.0059) (Fig. 6G,H). Our data suggest that, in combination with reduced pulmonary levels of S1PR1, the selective reduction of VEGFR-2 expression in lung contributes to the observed abnormalities in the pulmonary vasculature which is a feature characteristic of emphysema.

### Analysis of cytokines in BAL fluid showed an increase in OPN in Y^685^F-VE-cadherin mice

To identify the cytokine and chemokine profiles in the lungs, which are linked to changes in the local microenvironment, we analyzed the characteristics of the bronchoalveolar lavage fluid (BALF) of 4-week-old mice using a highly sensitive mouse angiogenic antibody array (Fig 6I). Out of the 53 detected proteins, only three (OPN, DLL4, IGFBP2) were found to be increased in KI mice, which supports no lung edema (See Extended Figures). Among these, only OPN had a significantly higher level in KI compared to WT (KI: 73.905 ± 17.24 vs WT: 44.95 ± 12.81; p-value=0.03; n=4 mice per genotype) (Fig. 6J). In addition its participation in the adhesion of ECs as described in our results above, OPN was shown to contribute to the expansion of the alveolar air space which characterized emphysema (42). Overall, the enhancement of barrier integrity by S1PR1 and the increase in airspace suggest that the lungs of KI mice may be analogous to emphysematous lungs in humans.

## Discussion

The VE-cadherin is the major structural component of adherens junctions in ECs and plays a key role in maintenance and regulation of endothelial barrier integrity. For the first time we provided new insights into lung EC gene expression in 4-week-old mice using our Y^685^F-VE cadherin knock-in mouse model. By investigating the effect of post-translational modification of the protein on gene expression, the introduction of a single mutation at Y^685^ to phenylalanine did not cause lethality, as the protein is an integral part of the structure of adherens junctions, in contrast to previous knockdowns of the VE-cadherin gene. As a result, it is important to note that 884 genes in the lung were altered, with 86.65% being downregulated and 13.34% being upregulated. It is worth highlighting that these changes in gene expression were specific to the lung, as there was no similar pattern observed in the kidney (FigEV7). This could be attributed to the inherent heterogeneity of ECs across different organs, which is closely linked to their distinct physiological functions.

Given that the ECs express Y685F-VE-cadherin, the transcriptomic analysis prompted us to examine the up-regulated and down-regulated genes separately in order to determine which processes were impaired or abnormally enhanced. Among the upregulated genes, four genes encode for G-protein-coupled transmembrane receptors including the Interleukin 9 receptor (IL9r), which belongs to the receptors of the hematopoietin superfamily (43), the S1PR1 G (Gαi/0) protein receptor for the signaling sphingolipid S1p (44). The third gene encodes for a G-protein-coupled receptor is GPR68, a receptor for sphingosylphosphorylcholine that shares a similar structure and function with ceramide, sphingosine-1-phosphate (S1P), and lysosphingolipids and was reported to induce the contraction of the vessel without [Ca^2+^]i elevation (45). In addition, the encoded protein is a proton-sensing receptor, inactive at pH 7.8 but active at pH 6.8 and capable of triggering intracellular signaling in response to alterations in extracellular pH around physiological values (45). The fourth gene encodes for a Gαi protein-coupled receptor, the CXCR3 chemokine receptor which is a member of the CXC chemokine receptor family. All those four genes encode G-protein coupled receptors that belong to the GPCR family, which responds to various stimuli and promotes different cellular functions. Among the downregulated genes, there are genes encoding for protein phosphatase *Ptprd*, which encodes for the receptor-type tyrosine-protein phosphatase delta involved in many genetic and epigenetic abnormalities such as gene amplification and DNA methylations as shown in a previous study by deactivation of PTPRD (46) . The genes encoding for enzymes involved in membrane co-transport include *Scl36a2 (*Solute Carrier Family 36 Member 2), which encodes for an electrogenic proton/amino acid that is involved in transmembrane movement of amino acids and derivatives (47). The gene *Scl38a5* encoded for protein that cotransports neutral amino acids and sodium ions, coupled to an H^+^ antiporter activity (48). The gene *Scn3a* encodes for a voltage-gated sodium channels which are transmembrane glycoprotein complexes (49). All of these genes were downregulated, which raises the interesting possibility of defects in membrane polarization of KI ECs. Furthermore, other genes that are downregulated, specifically those involved in lipid metabolism such as Lrat, Dpep1, and Scube2, may lead to additional modifications to the membranes caused by the Y^685^F-VE-cadherin mutation (50, 51, 52). Further studies are needed to better understand the impact of this single mutation on the expression and function of transmembrane proteins in endothelial cells and vascular dysfunction.

In an attempt to characterize the potential signaling pathways driven by the mutation, we analyzed the VEGF/VEGF-R2 pathway as it was shown that VE-cadherin was a major negative regulator of this pathway (53). When activated, VEGFR2 exhibit five major autophosphorylation sites which are Y^951^ lies in the kinase insert domain, Y^1054^ and Y^1059^ are in the kinase domain, whereas Y^1175^ (Y^1173^ in mouse) and Y^1214^ are in the C-terminal portion of the receptor (17–19). In KI mice, both VEGFR2 mRNA and protein levels were reduced, along with its phosphorylation on site Y^1173^. This single autophosphorylation event at Y^1173^ is crucial for the binding of PLCγ and activation of the PLCγ-PKC pathway, which plays a major role in signaling for migration and proliferation processes involved in angiogenesis (18). These results are consistent with our *in vitro* findings on isolated lung ECs from WT and KI mice. However, there was no significant difference in sites Y^1054^ and Y^1059^ required for full kinase activity between the two genotypes, indicating that VEGFR2 is fully activated, suggesting no negative effect of VE-cadherin or the mutant, unlike previously described (53).

Among the signaling partners coexisting with VE-cadherin, significant attention has been given to the tyrosine kinase c-Src. We have previously shown that c-Src associates with VE-cadherin, leading to phosphorylation of the junctional protein *in vivo* and *in vitro* and rapid dissociation of adherens junctions (8, 10, 24). The phosphorylation/activation of c-Src relies on the kinase CSK, which phosphorylates the Y^527^ in the tail domain of c-Src. In the KI mice there was a lesser association of CSK with c-Src than in the WT and therefore the level of activation of c-Src, as detected by pY^418^-Src, was higher in KI ECs than in WT ECs as well as its interaction with Y^685^F-VE-cadherin. Although we have observed increased pY^418^c-Src association with the mutant VE-cadherin, the specific mechanism of this interaction remains to be elucidated. It is unclear whether this is dependent on phosphorylation processes. One potential hypothesis is that the mutant is more phosphorylated at site Y^731^ than the WT. As phosphorylation is a covalent reaction, this could create an additional binding site for c-Src to VE-cadherin, which could explain the increased affinity for c-Src. Biological structural studies of the interaction between the cytoplasmic domain of VE-cadherin and c-Src, including amino acid sequence and measurements of the respective affinities, would be necessary to answer this question. However, it is noteworthy that the cytoplasmic domain of VE-cadherin contains at least one PXXP sequence (^673^PPRP^676^), which may allow VE-cadherin to associate with the SH3 domain of c-Src. This sequence may underlie the observed physical and functional links between c-Src and VE-cadherin in both WT and KI mice (54).

Cell-matrix or cell-cell junctions have been proposed to allow ECs to sense mechanical forces (55). In particular, our in vitro assays have shown that ECs have enhanced adhesion capabilities to FN and OPN, possibly through interactions with integrin receptors (24). Indeed, our transcriptomic analysis has shown a significant increase in the expression of the *ITGA5* gene in KI (3877.23, 6769.7, 1977.2) compared to WT (1275.78, 1068.35, 1295.35; p-value=0.00062). This gene encodes for α5β1, a member of the integrin family of ECM receptors which plays a crucial role in cell adhesion through a specific amino acid RGD sequence, which was initially identified as the binding site for FN. Further confirmation is required to determine whether differences in integrin protein expression are responsible for the distinct EC adhesion properties between KI and WT ECs (24). However, these results are consistent with changes observed in cell-cell junctions in KI. KI ECs had more VE-cadherin at the plasma membrane and were less dispersed under flow mimicking lung flow, suggesting greater resistance to mechanical forces. The lung is an organ with constant rhythmical changes in size that impinge ECs with rhythmical mechanical stresses so the importance of mechanical forces in alveolar function has the role of adherens junction and tight junction structures.

Among the different sites of phosphorylation of VE-cadherin, the Cheresh laboratory has highlighted sites Y^658^ and Y^731^ because they are binding sites for the interacting proteins p120 and β-catenin, respectively (34). However, a comparison of cadherin cytoplasmic domains shows that Y^731^ site is unique to VE-cadherin, suggesting that regulation of this residue may involve an endothelium specific mechanism. As a result, in our KI mouse model, we have demonstrated that it is the site Y^731^ which more phosphorylated *in vivo*. As a consequence, the association β-catenin-VE-cadherin was reduced in the KI mouse. Therefore, β-catenin translocated to the nucleus (35) and induced transcription by interacting with TCF/LEF (lymphoid enhancer factor) DNA-binding proteins to bind to the promoters of its target genes including FOXF1 (35, 38). FOXF1 has been shown to be abundant in the quiescent adult lung. Furthermore, FOXF1 expression decreased in ECs in human lung disease (38). Our results showed that increased binding of FOXF1 to the s1pr1 promoter increased S1PR1 mRNA and protein levels. Within the set of genes found to be upregulated in the KI mice, we focused on S1PR1 because VE-cadherin has previously been implicated as a target for S1P signaling in cultured ECs (56). In addition, S1PR1 has been described to inhibit VEGF-dependent vessel sprouting through a mechanism dependent on VE-cadherin function (56). Previous work showed that S1P which is the ligand of S1PR1 dramatically enhanced FGF-2-induced vascular density and the appearance of mature vascular structures during angiogenic processes (57). It is important to note that in our transcriptomic data the *fgfr4* gene, which is specific to FGFs 1, 2, 4, 6, 8, and 9, was down-regulated in Y^685^F-VE-cadherin (Log2Fold change -2.98, p-value= 9.99 x 10^-7^), as well as the *pdgfr* gene (Log2Fold change -2.42, p-value=2.38 x 10^-5^). The angiogenesis process is characterized by cross-talk between VEGF, FGF, and PDGF signaling pathways, as shown in numerous reports in the literature (58). We conclude that the observed decrease inn pulmonary vessel density in KI mice results from Y^685^F-VE-cadherin reducing the expression of three receptors (VEGFRs, FGFRs, PDGFRs) in pulmonary ECs and associated signaling pathways. The thin region of the blood-gas barrier with a thickness of only 0.2–0.3μm has only three layers: the capillary endothelium, the alveolar epithelium, and the ECM. The increased thickness of the lung vessels was observed in KI mice. It was shown that most of the thickening of basement membranes is associated with the capillary EC rather than the alveolar epithelial cell (59).This was seen in the pulmonary capillaries of patients with mitral stenosis and pulmonary venoocclusive disease (60). Therefore, the 4-week-old KI mice might be a good model for studying chronic obstructive pulmonary disease including emphysema since the severity of the disease was shown to depend upon endothelial dysfunction (61).

Our results demonstrated a decrease in collagen deposition in the lungs of KI mice which is consistent with the reduced expression of the gene *col13a1* as indicated by a Log2FC -3.64, (p-value = 2.62 x 10^-13^) in the transcriptomic data set. Previous analysis of type XIII collagen has shown that it is a non-fibrillar collagen that is cell-associated and plays a role in cell-matrix adhesion. Collagen XIII is found in practically all tissues studied, often in association with basement membrane or junctional structures where It also enables the transmission of mechanical signals between cells (62). In the developing lung, type XIII collagen is upregulated at the junctions between the parenchyma and the branching primary and secondary bronchioles, coinciding with the initiation of branching events (63). Further research is required to investigate the direct interaction between VE-cadherin and collagen XIII.

Another important extracellular matrix protein whose secretion was found to be increased in KI is OPN, which plays a critical role as a mediator in the pathogenesis of pulmonary vascular remodeling, cell attachment and during cyclic stretch (27). Its upregulation in KI mice during constant cyclic stretch suggests its potential involvement in the lung tissue response to mechanical stress, which may occur in chronic obstructive pulmonary disease (COPD) or acute respiratory distress syndrome (ARDS).

Lung morphology depends on the formation of a capillary network that regulates alveolar development. Physical forces resulting from repeated inflation/deflation can be translated into biological signals by several mechanisms. These include distortion of the cell membrane, which stimulates ion channels, and distortion of the cytoskeleton, which affects the nucleus and leads to changes in transcription. Our study has shown that mutation of a single tyrosine site in VE-cadherin affects specific cellular responses and modulates gene expression in the lung. Previous studies have reported knock-in mice in which the cytoplasmic tyrosine of a growth factor receptor has been substituted with phenylalanine, notably in the VEGF receptor (64). It is unusual for a single substitution to induce such changes in gene expression. Further research is therefore needed to understand how intercellular junctions based on Y^685^F-VE-cadherin affect the physiology of the pulmonary endothelium.

## Conclusion

In previous work we found that, in a highly angiogenic human glioblastoma, capillaries expressed pY^685^VE-cadherin, which was phosphorylated in a Src-dependent manner. This was associated with vascular structural abnormalities such as irregularity and tortuosity, suggesting abnormal endothelial cell-cell interactions and modified adherens junctions. Since VE-cadherin has been shown to contribute to endothelial differentiation and stability, we investigated whether the amino acid Y^685^ in the cytoplasmic domain of VE-cadherin was involved in an endothelial-specific transcriptomic program using a knock-in transgenic mouse model (Y^685^F-VE-cadherin). Our results demonstrate that a wide range of differentially expressed genes are found in the lung ECs of mice carrying the Y^685^F VE-cadherin mutation compared to WT, including the endothelial-specific gene encoding the S1PR1. We propose a novel mechanism by which mutant VE-cadherin is phosphorylated at site Y^731^, inducing translocation of b-catenin to the nucleus, activation by FoxF1 and transcription of the s1pr1 gene. This process has consequences for the lung vasculature as it leads to increased vessel wall thickness in vivo. Therefore, it may be envisioned that inhibiting VE-cadherin Y^685^ phosphorylation in cancer could redirect it to the amino acid 731, which is unique to VE-cadherin. Consequently, this would restore the thickness of tumor vessels, increase vascular normalization, and improve access of chemotherapy to the tumor cells. This hypothesis provides a basis for further research leading to the development of drugs that exclusively target the Y^685^ phosphorylation site of VE-cadherin without affecting other phosphosites. This would significantly increase the specificity of the drug by focusing on three key targets: 1) the vasculature, 2) VE-cadherin, and 3) the Y^685^ site in the LYAQV amino acid sequence of the cytoplasmic domain of VE-cadherin.

## Materials and Methods

### Animals

Four-week-old male wild-type (WT) C57BL/6 mice and four-week-old male transgenic Y^685^F-VE-cadherin (KI) C57BL/6 mice were used for the experiments and were generated as previously described (13). Mice were maintained in a conventional animal facility, on a 12-hour light/12-hour dark cycle. Food and water were available ad libitum. All procedures were carried out in compliance with the principles and guidelines established by the National Institute of Medical Research (INSERM) and approved by the Institutional by the Institutional Ethical Committee for animal experimentation (Comité d’Ethique).

### Reagents

Reagents were purchased from several source: collagenase type I (10114532, Fisher Scientific), DMEM (4.5g/L Glucose) (11965092, Gibco), EGM-2 BulletKit (CC-3156 & CC-4176, Lonza). Pierce BCA dosage Assay kit (23225, ThermoFisher). Protein G Sepharose 4B fast flow (P3296, Sigma Aldrich) was used for the Co-immunoprecipitation and ChIP assay. For RNA extraction Nucleospin® RNA (740955.50, Machery-Nagel) was used. For cDNA synthesis, 5x iScript Reaction Mix (1708890, Biorad) was used. For qPCR, SsoAdvanced Universal SYBR Green Supermix (1725270, Biorad) was purchased. For PCR, EconoTaq Plus Greeen (30033, Lucigen) was purchased. Mouse Angiogenesis array kit was purchased at ARY015, R&D system for protein array. For Masson trichrome staining, a special kit (H15) and hematoxylin Gill (GHS332) and eosin (HT110232) were purchased from Sigma Aldrich.

### Antibodies

Commercially available antibodies were purchased from several sources: Antibodies used for the FACS: CD-31 (PE) antibodies were from Becton-Dickinson, CD45R (53-0452-82, Fisher scientific). For Miltenyi cell isolation: CD45 microbeads (130-052-301, Miltenyi), CD31 microbeads (130-097-418, Miltenyi), MS Separate column (130-042-201, Miltenyi). Anti-VE-cadherin antibody (AF1002) and anti FOXF1 antibody (AF4798) were purchased from R&D system. Anti S1PR1 antibody was from Sigma-Aldrich (SAB4500687, Anti-VEGFR2 antibody (sc-6251), anti-VEGF antibody (sc-7269), anti-Src antibody (sc-18), anti-VE-cadherin antibody (sc-19) were from Santa-Cruz. Anti-pY^1175^-VEGFR2 antibody (D5B11), and anti β-catenin antibody (9562) were purchased from Cell signaling. Anti-pY^418^-Src (LF-PA20465) was from Ab Frontier. Anti-pY^658^-VE-cadherin antibody (44-1144G), and anti-pY^731^-VE-cadherin antibody (44-1145G) were from Invitrogen. Cy3 Donkey anti-Goat (705-166-147) secondary antibody was from Jackson.

### Quantitative real time RT-PCR (qRT-PCR)

#### mRNA isolation from the endothelial cells

Nucleospin® RNA kit was used for the isolation of RNA from the cultured cells. The protocol was followed exactly according to the manufacturer. RNA was eluted with 40µL ofautoclaved distilled water The purity of RNA was checked and quantified with a NanoDrop spectrophotometer (Thermo Fisher, Massachusetts, USA)

#### Reverse transcription of RNA to cDNA

The conversion of RNA isolated from the ECs to cDNA was done using iScript cDNA Synthesis Kit. The sample reaction was prepared following the manufacturer instructions. The complete reaction mix was incubated in a thermal cycler in the following protocol: 5 minutes at 25°C, 20 minutes at 46°C and 1 minute at 95°C.

#### Quantitative Polymerase Chain Reaction (qPCR)

SsoAdvanced Universal SYBR Green Supermix was used for cDNA amplification with different genes primers (see FigEV 10) on the CFX96 Real-Time PCR Detection System using the following conditions: 95°C 30 seconds/ 95°C 5 seconds; 60 °C 15 seconds for 39 cycles, 95°C for 10 seconds, 65°C for 5 seconds and 65°c for 5 seconds.

#### Polymerase Chain Reaction PCR

DNA samples from ChIP were analyzed by qPCR. DNA were amplified with FOXF1 binding promoter primers (see table S1) and EconoTaq Plus Greeen according the following cycles: 3 minutes at 95°C, 34 cycles: 95°C 15 seconds; 62°C 15 seconds; 72°C 20 seconds, 72°C 10 minutes. The samples were loaded onto the agarose gel for analysis.The relative level of RNA was normalized to housekeeping gene *hprt1*. Relative gene expressions were calculated using the DCT method (DCT= DCT treated-DCT untreated, normalised to the control) and fold change was calculated: linearized DCT treated/ linearized DCT control (65).

#### Transcriptomic analysis

The ECs from the lung was isolated with the FACS method as described before and the cell pellet were frozen in nitrogen liquid and sent to GENEWIZ for RNA-seq (Genomics European Headquarters Bahnhofstr. 8 04158 Leipzig Germany).

#### RNA Extraction, Library Preparation, NovaSeq Sequencing, and Standard RNA-Seq analysis

Total RNA was extracted using Qiagen RNeasy Mini kit following manufacturer’s instructions (Qiagen, Hilden, Germany). RNA samples were quantified using Qubit 4.0 Fluorometer (Life Technologies, Carlsbad, CA, USA) and RNA integrity was checked with RNA Kit on Agilent 5300.

RNA sequencing libraries were prepared using the NEBNext Ultra RNA Library Prep Kit for Illumina following manufacturer’s instructions (NEB, Ipswich, MA, USA). Briefly, mRNAs were first enriched with Oligo(dT) beads. Enriched mRNAs were fragmented for 15 minutes at 94 °C. First strand and second strand cDNAs were subsequently synthesized. cDNA fragments were end repaired and adenylated at 3’ends, and universal adapters were ligated to cDNA fragments, followed by index addition and library enrichment by limited-cycle PCR. Sequencing libraries were validated using NGS Kit on the Agilent 5300 Fragment Analyzer (Agilent Technologies, Palo Alto, CA, USA), and quantified by using Qubit 4.0 Fluorometer (Invitrogen, Carlsbad, CA, USA).

The sequencing libraries were multiplexed and loaded on the flowcell on the Illumina NovaSeq 6000 instrument according to manufacturer’s instructions. The samples were sequenced using a 2x150 Pair-End (PE) configuration v1.5. Image analysis and base calling were conducted by the NovaSeq Control Software (NCS). Raw sequence data (.bcl files) generated from Illumina NovaSeq was converted into fastq files and de-multiplexed using Illumina bcl2fastq program version 2.20. One mismatch was allowed for index sequence identification.

After investigating the quality of the raw data, sequence reads were trimmed to remove possible adapter sequences and nucleotides with poor quality using Trimmomatic v.0.36. The trimmed reads were mapped to the mouse reference genome available on ENSEMBL using the STAR aligner v.2.5.2b. BAM files were generated as a result of this step. Unique gene hit counts were calculated by using feature Counts from the Subread package v.1.5.2. Only unique reads that fell within exon regions were counted.

After extraction of gene hit counts, the gene hit counts table was used for downstream differential expression analysis. Using DESeq2, a comparison of gene expression between the groups of samples was performed. The Wald test was used to generate P values and Log2 fold changes. Genes with adjusted P values < 0.05 and absolute log2 fold changes >1 were called as differentially expressed genes for each comparison. Gene ontology analysis was performed on the statistically significant set of genes by implementing the software GeneSCF. The goa_Mus musculus GO list was used to cluster the set of genes based on their biological process and determine their statistical significance.

A PCA analysis was performed using the “plotPCA” function within the DESeq2 R package. The plot shows the samples in a 2D plane spanned by their first two principal components. The top 500 genes, selected by highest row variance, were used to generate the plot.

The functional enrichment analysis was performed based on the “Biological Process” ontologies from the Gene Ontology database (GO.db R package version 3.16.0). ORA and GSEA were computed using the Bioconductor/R package clusterProfiler (version 4.6.0). Gene sets with a corrected p-value (Benjamini-Hochberg correction method) < 0.05 were considered significant. Protein-protein networks were built using the STRING-DB web software (https://string-db.org/) using significant DEGs found in cell-cell adhesion or angiogenesis ontologies. Statistical analysis were performed using R (version 4.2.2).

#### Cell migration assay

The freshly isolated ECs were immediately seeded at 200 000 cells / well in 96 well plates pre-coated with fibronectin and grown to confluency in the EGM-2 medium. The day of the wound assay, the cells were washed with PBS (Ca+/Mg+) and the scratch was performed with the Incucyte® Cell Migration Kit (Sartorius). The cells were washed carefully twice with PBS (Ca+/Mg+) then DMEM (4.5G/L Glucose) containing 1.5% FBS was added. The full time course migration was performed for 20h in the Incucyte Zoom live-cell analysis system (Sartorius) and the analysis of the gap area of the wound was performed by the Incucyte Basic Analyzer software.

#### Proliferation assay

Isolated ECs were seeded onto 96 well-plates pre-coated with fibronectin in EGM-2 and the proliferation assay was performed for 60 hours in the Incucyte proliferation assay according to (20) (Sartorius). The analysis of the cell proliferation was performed by the Incucyte Basic Analyzer software.

#### 3D fibrin beads gel assay in vitro

3D fibrin gel bead assay was performed as described previously (22). Isolated ECs were mixed with Cytodex 3 microcarrier beads at the concentration of 800 ECs per beads. In 1 mL of warm EGM-2 medium. Beads and ECs were then co-incubated in a humidified incubator at 37 °C and 5% CO2 and gently manually shaken every 20 min for 4 h to allow cell adherence to the bead surface. After 4 hours, they were transferred into 25cm² flask in 5 mL of EGM-2 medium overnight. The following day, the coated beads were resuspended in 2.5mg/ml fibrinogen solution, 1µg/mL aprotinin and 10ng/mL VEGF at concentration of ̴ 500 beads/ml. 12µL of thrombin 10U/mL and 288µL of the mix fibrinogen/beads suspension per well of 24-well plate. The plate was incubated 15 minutes in a humidified incubator at 37 °C and 5% CO2 until clot formation. 500µL of EGM-2 supplemented with angiopoetin 1 (250µg/mL), hGF (100ng/mL), IGFBP3 (50ng/mL), VEGF (20ng/mL), PDGF (10ng/mL), EGF (100ng/mL) (20). After 4 days of incubation, the angiogenic sprouting were imaged with the Axioobserver Z1 Zeiss software with objective 5X/0.15 Numerical Aperture. Quantification of morphometrical parameters of the capillary network was performed using a program developed for the ImageJ software (22). This plugin is an extension of the “Angiogenesis Analyzer” for Image J written in the macro language of ImageJ. Representative parameters measured were number of anchorage per junctions, number of junctions, total master segments and number of meshes per beads.

#### Adhesion assay

Isolated ECs were seeded in 96 well-plates pre-coated with fibronectin (10 μg/mL) or OPN (0.3 μg/mL) in EGM-2 (1500 cells/well). After 30 min of incubation, the coverslip-attached cells were washed with PBS (Ca+/Mg+), fixed with PFA 4% and permeabilized with 0.2% Triton X100 for 20 minutes at room temperature. The nuclei were stained with Hoescht (1/400) for 5 minutes in PBS. After 3 washes with PBS (Ca+/Mg+), the cells were imaged with the Axio-observer Z1 Zeiss software with objective X5/0.15 Numerical Aperture. The number of nuclei were counted with Image-J software (NIH, Bethesda, MD).

#### Immunofluorescence

Isolated ECs were cultured 4-7 days with EBM-2 and then washed with PBS (Ca+/Mg+). The cells were fixed with PFA 4% and permeabilized Triton 0.2% 20 minutes at room temperature. The cells were washed twice for 5 minutes with PBS (Ca+/Mg+) and blocked with PBS (Ca+/Mg+) / BSA 3% for 30 minutes at room temperature. The cells were incubated with the mouse VE-cadherin primary antibody (2μg/mL) in PBS (Ca+/Mg+) / BSA 3% for 1h at room temperature with gentle agitation. After two washes with PBS (Ca+/Mg+), Tween 0.1% for 5 minutes, Cy3 Donkey anti-Goat antibody (1/200) was added in the dark 1h at room temperature under gentle agitation. The nuclei were stained with Hoechst in PBS for 5 minutes at room temperature in the dark before the slides were mounted. The slides were imaged with the Axio Imager M2 Zeiss microscope (with objective X63/ Numerical Aperture 1.4, oil immersion). VE-cadherin staining was quantified by measuring the staining area at the membrane and the integrated density of the staining using ImageJ software macro (NIH, Bethesda, MD). Scale bars are 20 μm.

#### Analysis of VE-cadherin Immunostaining

VE-cadherin signal and nuclei were segmented by first denoising images with a Gaussian filter (σ = 1), and then denoised images were threshold with the triangle algorithm (65). The mean of VE-cadherin signal was computed over its segmentation mask, and the total area of nuclei was computed from Hoescht channel mask. In order to obtain an invariant measure of VE-cadherin signal regarding to both cell number and cell size, the mean VE-cadherin was divided by the total area of nuclei.

#### Flow-Induced Orientation Analysis

To analyze the orientation of cells, we calculated the angle formed between the vector of the flow direction (obtained by knowledge of flow direction within the slide) and the “orientation vector” given by orientation of the main axe of each cell. Each value for each cell is then used to plot the hemi-roses presented in Figure 4K. To plot the bar graphs presented in Figures 4E K, the cells were classified in 2 categories: aligned with the flow direction, and not aligned with the flow direction. For in vitro experiment, aligned with the flow was defined as an absolute value of angle between 0 and 45, not aligned with the flow as an absolute value of angle between 45 and 90; Angle calculation and roses presentation was done automatically using a homemade Matlab script validated on the first experiment by a comparison to hand calculation with FIJI software (66).

#### Chromatin immunoprecipitation assay (ChIP)

The ChIP protocol was kindly given by Dr Marco Saponaro. Briefly, 4 week-old male mice WT and KI was sacrificed and the lungs were harvested and placed immediately in cold PBS (Ca-/Mg-). Immersed tissues were shredded using scalpel into small pieces and digested with 4 mL of collagenase I (3mg/mL) in DMEM (4.5g/L Glucose) with no serum at 37°C for 45 min. Every 15 minutes, the tube was shaken to improve the collagenase I activity. The digestion is stopped with 20% FBS. The cells suspension was pass through a 10 mL syringe with a 18G needle and through several filters: 100, 70µm filters followed by centrifugation at 230 x g for 5 minutes. The supernatant was carefully removed and resuspended in PBS. Formaldehyde was added to 1 % final concentration and incubated in rotation at room temperature for 15 minutes. Glycine (125mM) was immediately added and followed by 5 minutes incubation at room temperature under agitation. The solution was centrifuged 2 minutes at 230 x g and washed twice with ice-cold PBS for 2 minutes at 230 x g. Cells were lysed with ChIP buffer (5mM HEPES, pH 8.0, 85mM KCL, 0.5% NP-40 and leupeptin (10µg/mL), Na3VO4 (2mM)), and incubated 5 minutes on ice. The cell solution was centrifuged 5 minutes at 3900 x g at 4°C to obtain the nuclei pellet. The nuclei were lysed with ChIP nuclear lysis buffer (50mM Tris/HCl pH 8.1, 10mM EDTA, 1% SDS + leupeptin (10µg/mL) and Na3VO4 (2mM)) and incubated 5 minutes on ice. Nuclear lysate was sheared by using an ice-water bath embedded sonication system at high power, 30 seconds on, 30 seconds off mode for 10 minutes. Sonicated chromatin was cleared by centrifugation at 20,000 g for 10 min at 4°C. Chromatin was diluted 1:5 with ChIP dilution buffer (0.01% SDS, 1.1% Triton X-100, 1.2mM EDTA (pH 8.0), 16.7mM Tris/HCl pH 8.1, 167mM NaCl, leupeptin (10µg/mL), Na3VO4 (2mM)). The sample was cleared with 30µL of Sepharose beads G for 30 minutes under agitation at 4°C and centrifugation at 3,800 x g 2 minutes at 4°C. The supernatant was transferred into a new tube and 2.5µg of FOXF1 antibody was added overnight at 4°C under agitation. Protein G Sepharose beads were added into the tubes and incubated 3h at 4°C under agitation. Washes are performed sequentially, doing them two times with 1ml of each of the following buffers: ChIP low salt buffer (0.1% SDS, 1% Triton X-100, 2mM EDTA, 20mM Tris/HCl pH 8.1, 150mM NaCl) and ChIP high salt buffer (0.1% SDS, 1% Triton X-100, 2mM EDTA, 20mM Tris-HCl pH 8.1, 500mM NaCl) followed by ChIP LiCl buffer (10mM Tris-HCl pH 8.0, 250mM LiCl, 1%NP40, 1% deoxycholic acid, 1mM EDTA). A final wash was made with TE buffer (pH=8) and centrifuged at 1,000 x g for 3 minutes. The pellet was resuspended in 40μL elution buffer (50mM Tris/HCL pH 8.0, 10mM EDTA, 1%SDS) with RNAse 10mg/L and incubated overnight at 65°C. The sample was centrifuged 1 minute at 12000 rpm and the supernatant was transferred into a fresh tube where 120µL TE/1%SDS was added with proteinase K (20mg/mL) and glycogen (10mg/mL) whereas the pellet were resuspended in Laemmli 2X for western blot analysis. The chromatin solution was incubated for 2 hours at 37°C. An exact equal volume of Phenol/Chloroform/Isoamilalchol 25:24:1 pH 8.0 was added and mixed by inverting the tube until the phases are completely mixed. Then, the tube was centrifuged at 15,000 x g 5 minutes and the upper layer was collected into a new tube where ethanol 100% and 5M NaCl were added. The sample was stored at -20°C for 1 hour. It was centrifuged again 10 minutes at 15,000 x g . The pellet was resuspended in ethanol 70 % and centrifuged 10 minutes 15,000 x g . The supernatant was retrieved as the pellet let completely dry before adding TE buffer to perform the PCR.

#### Histological sectioning of lungs and staining

For all histological procedures, the lungs were inflated to full capacity and fixed by intratracheal instillation of formalin 10% (1 mL), immersed in formalin overnight, and then embedded in paraffin. Briefly, mice were sacrificed and lung were harvested quickly and transferred in ethanol 90% 2 hours at room temperature, then in ethanol 100% 3 hours still at room temperature. After 4 times in methylcyclohexane for 45 minutes at room temperature, the lungs were placed in paraffin/methylcyclohexane (1:1) at 60°C for 24 hours. Lungs were transferred into fresh paraffin every day for 4 days, and let to dry at room temperature. Paraffin embedded lungs were sagittally sectioned at 5 μm and kept at 60°C for an hour. Sections were stained with Harris’s hematoxylin/eosin or Masson’s trichrome stain for general morphology and morphometric analysis. Images post analysis was performed with Image-J software (NIH, Bethesda, MD).

#### Pulmonary vessel quantification

Pulmonary arteries from 4 week-old mice (4 WT and 4 KI) were analyzed. Measurements of the vessel number were taken for each slide corresponding to each mouse lung. Only arteries between 50-150µm of luminal diameter were selected for analysis. We used Image J software for measurement of total vessel area (µm²), luminal area (µm²), and inner circumference (µm). Medial area (µm2) was calculated as the difference between total and luminal areas. Standard medial thickness (µm) was calculated as the ratio of medial area to inner circumference (67).

#### Lung morphological assessment

After fixation, 5-μm sections were stained with hematoxylin and eosin. The mean linear intercept, as a measure of interalveolar wall distance, was determined by light microscopywith objective ×100. The mean linear intercept was obtained by dividing the total length of a line drawn across the lung section by the total number of intercepts encountered in 72 lines per each rat lung, as previously described (67).

#### Sub-cellular EC fractionation

ECs were lysed in 5mM HEPES/NaOH pH 8.0, 85mM KCL, 0.5% NP-40 and leupeptin (10µg/mL), Na3VO4 (2mM) lysis buffer, and incubated 10 minutes on ice. The cell solution was centrifuged for 5 min at 3900 g at 4°C and the pellet was lysed with nuclear lysis buffer (50mM Tris/HCL pH 8.1, 300mM NaCl, 10mM EDTA, 1% SDS, leupeptin (10µg/mL), Na3VO4 (2mM)) and incubated 10 minutes on ice. Nuclear extracts were obtained by sonication (30 seconds on, 30 seconds off mode for 10 minutes) followed by centrifugation at 20,000xg for 10 min at 4°C. The supernatant was collected as the nuclei fraction.

#### Western blot analysis and immunoprecipitation

Protein extracts from lung and isolated ECs were analyzed by Western blot analysis as previously described^8^. The antibobies used in the present study were: VE-cadherin (AF1002) (1/500), pY^731^VE-cadherin (1/500), pY^658^VE-cadherin (1/500), pY^416^Src (1/1000), Src (1/500), β-catenin (1/1000), VEGFR2 (1/500), pY^1173^VEGFR2 (1/500), VEGFA (1/500), S1PR1 (1/250). Protein expression was normalized with actin or vinculin expression level as indicated. The results are presented as mean ± SEM.

For immunoprecipitation, 100μg of proteins were incubated with 1µg of VE-cadherin antibody (sc-19) overnight. Then the samples were incubated with protein G Sepharose for 3 hours, then washed run onto SDS-PAGE 10%. The proteins were then transferred onto a 0.45μm nitrocellulose membrane. Membranes were blocked with milk, incubated with primary antibodies and corresponding HRP-conjugated secondary antibodies. Immunoreactive proteins were visualized by Bio-Rad ChemiDoc™ with ECL reagent (Bio-Rad Laboratories) and were quantified using densitometry with Image-J software (NIH, Bethesda, MD).

#### Broncho-alveolar lavage (BAL)

Using a 5-mL syringe, 1 mL of PBS with protease and phosphatase inhibitor was injected slowly into a cannula connected to the mice larynx. The BALF was collected by slowly drawing the injected PBS back into the syringe and was place in a 2-mL microfuge tube and place it on ice until BAL samples from all mice are collected. When all were collected, after a centrifugation for 5 min at 1000g, the supernatants were collected and concentrated on Amicon Ultra-2 - Ultracel-10 membrane, 10 kDa. Protein concentrations was determined using the Micro BCA Protein Assay Kit. Analysis was performed using the Angiogenesis assay according the manufacturer’s instructions (4 membranes). Immunoreactive proteins were visualized by Bio-Rad ChemiDoc™ with ECL reagent (Bio-Rad Laboratories) and were quantified using densitometry with Image-J software (NIH, Bethesda, MD).

#### Angiogenesis Assay

Analysis of the protein expression in the broncho-alveolar lavage of mice was performed with the Mouse Angiogenesis Array Kit. Briefly, the BAL from 4 week-old male mice of each genotype was centrifuged at 1000g and the supernatant was concentrated on Amicon Ultra-2 - Ultracel-10 membrane, 10 kDa and protein concentrations was determined using the Micro BCA Protein Assay Kit. The same amount of protein concentration for each mouse genotype was analyzed using the Angiogenesis assay according the manufacturer instructions (4 membranes). Immunoreactive proteins were visualized by Bio-Rad ChemiDoc™ with ECL reagent (Bio-Rad Laboratories) and were quantified using densitometry with Image-J software (NIH, Bethesda, MD).

#### Statistical analysis

At least three to five mice per group were used in each set of experiments. The experiments were performed at least three times under identical conditions with comparable results. Data are represented as means ± SD. The sample size, what each “n” represents, the statistical tests used and the result of the statistical test are indicated in each respective figure legend. Data was analyzed using the software GraphPad Prism version v8.0 (GraphPad Software Inc., La Jolla, CA, USA) or R software as indicated in each respective figure legend. If data deviated from a normal distribution, and/or displayed variance heterogeneity, statistical analysis were performed on log-transformed data or with the use of non-parametric test. Within each experiment, differences in means between groups for each parameter were analysed using either a one-way ANOVA or Kruskal-Wallis test with Tukey’s post hoc test or the non-parametric Wilcoxon rank sum test. Differences between two different groups were analyzed by unpaired Student’s t-test. A probability value <0.05 indicated by an asterisk (*) showed a significant difference between 2 groups. Non significant difference was indicated by ns. On all graphs data are presented as mean ± SEM, and in all analyses *P-*values <0.05 were considered statistically significant. For immunostaining and immunofluorescence images, assessment of the differences was performed using boxplot and scatter plot in R along with Kruskal-Wallis rank sum test and chi-square test to determine the p-value.

## Acknowledgments

The authors are indebted to: Soumalamya Bama-Toupet, Charlène Magallon and Hervé Pointu for their technical assistance in the animal facilities, Aude Salomon and Violaine Simon for their excellent technical expertise, Véronique Colin-Faure as the IRIG flow cytometry platform manager and we thank her for excellent technical support for cell sorting, Frédéric Sergent and Nicolas Lemaitre for their technical expertise regarding immunohistochemistry of lungs, Frédérique Mitler and Frédérique Kermarrec-Marcel for their advices in immunoblot and PCR, Patricia Obeid for her expertise in image quantifications and all members of the BIOMICS team for their support. We thank a lot Mylène Pezet from the Optimal Platform, and Dr Eva Faurobert for their expertise in primary cell culture. We thank Dr Saporano for excellent technical advices for CHIP assays.

## References

1. Aird WC. Phenotypic heterogeneity of the endothelium: I. Structure, function, and mechanisms. Circ Res. 2007;100(2):158–73. doi: 10.1161/01.RES.0000255691.76142.

2. Grunewald M, Avraham I, Dor Y, Bachar-Lustig E, Itin A, Jung S, Chimenti S, Landsman L, Abramovitch R, Keshet E. VEGF-induced adult neovascularization: recruitment, retention, and role of accessory cells. Cell. 2006, 124(1):175–189.doi: 10.1016/j.cell.2005.10.036.

3. Rafii S, Butler JM, Ding BS. Angiocrine functions of organ-specific endothelial cells. Nature. 2016, 529:316–325. doi: 10.1038/nature17040.

4. Ferrara N, Houck KA, Jakeman LB, Winer J, Leung DW. The vascular endothelial growth factor family of polypeptides. J Cell Biochem. 1991, 47(3):211–218. doi: 10.1002/jcb.240470305.

5. Breviario F, Caveda L, Corada M, Martin-Padura I, Navarro P, Golay J, Introna M, Gulino D, Lampugnani MG, Dejana E. Functional properties of human vascular endothelial cadherin (7B4/cadherin-5), an endothelium-specific cadherin. Arterioscler Thromb Vasc Biol. 1995, (8):1229–1239. doi: 10.1161/01.atv.15.8.1229.

6. Kouklis P, Konstantoulaki M, Malik AB. VE-cadherin-induced Cdc42 signaling regulates formation of membrane protrusions in endothelial cells. J Biol Chem. 2003. 278: 16230–16236. doi: 10.1074/jbc.M212591200.

7. Esser S, Lampugnani, MG, Corada M, Dejana E, Risau W J. Cell Sci. 1998;111, 1853–1865. doi: 10.1242/jcs.111.13.1853.

8. Lambeng N, Wallez Y, Rampon C, Cand F, Christé G, Gulino-Debrac D, Vilgrain I, Huber P. Vascular endothelial-cadherin tyrosine phosphorylation in angiogenic and quiescent adult tissues. Circ Res. 2005 ;96(3):384–391. doi: 10.1161/01.RES.0000156652.99586.

9. Lindquist J, Simeoni L, Schraven B. Transmembrane adapters: attractants for cytoplasmic effectors. Immunol Rev. 2003, 191: 165–182. doi: 10.1034/j.1600-065x.2003.00007.

10. Wallez Y, Cand F, Cruzalegui F, Wernstedt C, Souchelnytskyi S, Vilgrain I, Huber P. Src kinase phosphorylates vascular endothelial-cadherin in response to vascular endothelial growth factor: identification of tyrosine 685 as the unique target site. Oncogene 2007, 26(7):1067–77. doi: 10.1038/sj.onc.1209855.

11. Das S, Marsden PA. Angiogenesis in glioblastoma. N. Engl. J. Med. 2013;369:1561–1563. doi: 10.1056/NEJMcibr1309402.

12. Vilgrain I, Sidibé A, Polena H, Cand F, Mannic T, Arboleas M, Boccard S, Baudet A, Gulino-Debrac D, Bouillet L, Quesada JL, Mendoza C, Lebas JF, Pelletier L, Berger F. Evidence for post-translational processing of vascular endothelial (VE)-cadherin in brain tumors: towards a candidate biomarker. PLoS One. 8(12):e80056. (2013) doi: 10.1371/journal.pone.0080056.

13. Sidibé A, Polena H, Pernet-Gallay K, Razanajatovo J, Mannic T, Chaumontel N, Bama S, Maréchal I, Huber P, Gulino-Debrac D, Bouillet L, Vilgrain I. VE-cadherin Y685F knock-in mouse is sensitive to vascular permeability in recurrent angiogenic organs. Am J Physiol Heart Circ Physiol. 2014;307(3):H455–63. doi: 10.1152/ajpheart.00774.2013.

14. Sidibé A, Polena H, Razanajatovo J, Mannic T, Chaumontel N, Bama S, Maréchal I, Huber P, Gulino-Debrac D, Bouillet L, Vilgrain I. Dynamic phosphorylation of VE-cadherin Y685 throughout mouse estrous cycle in ovary and uterus. Am J Physiol Heart Circ Physiol. 2014;307(3):H448–54. doi: 10.1152/ajpheart.00773.2013.

15. Wessel F, Winderlich M, Holm M, Frye M, Rivera-Galdos R, Vockel M, Linnepe R, Ipe U, Stadtmann A, Zarbock A, Nottebaum AF, Vestweber D. Leukocyte extravasation and vascular permeability are each controlled in vivo by different tyrosine residues of VE-cadherin. Nat Immunol. 2014;15(3):223–230. doi: 10.1038/ni.2824.

16. Morini MF, Giampietro C, Corada M, Pisati F, Lavarone E, Cunha SI, Conze LL, O’Reilly N, Joshi D, Kjaer S, George R, Nye E, Ma A, Jin J, Mitter R, Lupia M, Cavallaro U, Pasini D, Calado DP, Dejana E, Taddei A. VE-Cadherin-Mediated Epigenetic Regulation of Endothelial Gene Expression. Circ Res. 2018 Jan 19;122(2):231–245. doi: 10.1161/CIRCRESAHA.117.312392.

17. Kendall RL, Rutledge RZ, Mao X, Tebben AJ, Hungate RW, Thomas KA. Vascular endothelial growth factor receptor KDR tyrosine kinase activity is increased by autophosphorylation of two activation loop tyrosine residues. J Biol Chem. 1999;274(10):6453–60. doi: 10.1074/jbc.274.10.6453.

18. Dougher M, Terman BI. Autophosphorylation of KDR in the kinase domain is required for maximal VEGF-stimulated kinase activity and receptor internalization. Oncogene. 1999;18(8):1619–27. doi: 10.1038/sj.onc.1202478.

19. Takahashi T, Yamaguchi S, Chida K, Shibuya M. A single autophosphorylation site on KDR/Flk-1 is essential for VEGF-A-dependent activation of PLC-gamma and DNA synthesis in vascular endothelial cells. EMBO J. 2001 Jun 1;20(11):2768–78. doi: 10.1093/emboj/20.11.2768.

20. Nowak-Sliwinska P, Alitalo K, Allen E, Anisimov A, Aplin AC, Auerbach R, Augustin HG, Bates DO, van Beijnum JR, Bender RHF, Bergers G, Bikfalvi A, Bischoff J, Böck BC, Brooks PC, Bussolino F, Cakir B, Carmeliet P, Castranova D, Cimpean AM, Cleaver O, Coukos G, Davis GE, De Palma M, Dimberg A, Dings RPM, Djonov V, Dudley AC, Dufton NP, Fendt SM, Ferrara N, Fruttiger M, Fukumura D, Ghesquière B, Gong Y, Griffin RJ, Harris AL, Hughes CCW, Hultgren NW, Iruela-Arispe ML, Irving M, Jain RK, Kalluri R, Kalucka J, Kerbel RS, Kitajewski J, Klaassen I, Kleinmann HK, Koolwijk P, Kuczynski E, Kwak BR, Marien K, Melero-Martin JM, Munn LL, Nicosia RF, Noel A, Nurro J, Olsson AK, Petrova TV, Pietras K, Pili R, Pollard JW, Post MJ, Quax PHA, Rabinovich GA, Raica M, Randi AM, Ribatti D, Ruegg C, Schlingemann RO, Schulte-Merker S, Smith LEH, Song JW, Stacker SA, Stalin J, Stratman AN, Van de Velde M, van Hinsbergh VWM, Vermeulen PB, Waltenberger J, Weinstein BM, Xin H, Yetkin-Arik B, Yla-Herttuala S, Yoder MC, Griffioen AW. Consensus guidelines for the use and interpretation of angiogenesis assays. Angiogenesis. 2018. 21(3):425–532. doi: 10.1007/s10456-018-9613.

21. Sakurai Y, Ohgimoto K, Kataoka Y, Yoshida N, Shibuya M. Essential role of Flk-1 (VEGF receptor 2) tyrosine residue 1173 in vasculogenesis in mice. Proc Natl Acad Sci U S A. 2005;102(4):1076–81. doi: 10.1073/pnas.0404984102.

22. Carpentier G, Berndt S, Ferratge S, Rasband W, Cuendet M, Uzan G, Albanese P. Angiogenesis Analyzer for ImageJ - A comparative morphometric analysis of “Endothelial Tube Formation Assay” and “Fibrin Bead Assay”. Sci Rep. 2020;10(1):11568. doi: 10.1038/s41598-020-67289-8.

23. Levinson NM, Seeliger MA, Cole PA, Kuriyan J. Structural basis for the recognition of c-Src by its inactivator Csk. Cell. 2008;134(1):124–34. doi: 10.1016/j.cell.2008.05.051.

24. Vielreicher M, Harms G, Butt E, Walter U, Obergfell A. Dynamic interaction between Src and C-terminal Src kinase in integrin alphaIIbbeta3-mediated signaling to the cytoskeleton. J Biol Chem. 2007;282(46):33623–33631. doi: 10.1074/jbc.M704107200.

25. Nakagawa T, Tanaka S, Suzuki H, Takayanagi H, Miyazaki T, Nakamura K, Tsuruo T. Overexpression of the csk gene suppresses tumor metastasis in vivo. Int J Cancer. 2000;88(3):384–91. PMID: 11054667

26. Huebsch JC, McCarthy JB, Diglio CA, Mooradian, D L. Endothelial Cell Interactions With Synthetic Peptides From the Carboxyl-Terminal Heparin-Binding Domains of Fibronectin. Circ. Res. 1995; 77, 43–53.doi: 10.1161/01.res.77.1.43.

27. Barkas GI, Kotsiou OS. The Role of Osteopontin in Respiratory Health and Disease. J Pers Med. 2023;13(8):1259. doi: 10.3390/jpm13081259.

28. Clark RA, Folkvord JM, Nielsen LD. Either exogenous or endogenous fibronectin can promote adherence of human endothelial cells. J Cell Sci. 1986;82:263–280.doi: 10.1242/jcs.82.1.263.

29. Volberg T, Romer L, Zamir E, Geiger B. pp60(c-src) and related tyrosine kinases: a role in the assembly and reorganization of matrix adhesions. J Cell Sci. 2001;114(Pt 12):2279–89. doi: 10.1242/jcs.114.12.2279.

30. Roux E, Bougaran P, Dufourcq P, Couffinhal T. Fluid Shear Stress Sensing by the Endothelial Layer. Front Physiol. 2020;11:861. doi: 10.3389/fphys.2020.00861.

31. Filipovic N, Ghimire K, Saveljic I, Milosevic Z, Ruegg C. Computational modeling of shear forces and experimental validation of endothelial cell responses in an orbital well shaker system. Computer Methods in Biomechanics and Biomedical Engineering, 2016; 19: 581–590.

32. Bachir AI, Horwitz AR, Nelson WJ, Bianchini JM. Actin-Based Adhesion Modules Mediate Cell Interactions with the Extracellular Matrix and Neighboring Cells. Cold Spring Harb Perspect Biol. 2017;9(7):a023234. doi: 10.1101/cshperspect.a023234.

33. Mahabeleshwar GH, Feng W, Reddy K, Plow EF, Byzova TV. Mechanisms of integrin-vascular endothelial growth factor receptor cross-activation in angiogenesis. Circ Res. 2007;101(6):570–80. doi: 10.1161/CIRCRESAHA.107.155655.

34. Potter MD, Barbero S, Cheresh DA. Tyrosine phosphorylation of VE-cadherin prevents binding of p120- and beta-catenin and maintains the cellular mesenchymal state. J Biol Chem. 2005;280(36):31906–31912. doi: 10.1074/jbc.M505568200.

35. McCrea PD, Gottardi CJ. Beyond β-catenin: prospects for a larger catenin network in the nucleus. Nat Rev Mol Cell Biol. 2016;17(1):55–64. doi: 10.1038/nrm.2015.3.

36. Allende ML, Yamashita T, Proia RL. G-protein-coupled receptor S1P1 acts within endothelial cells to regulate vascular maturation. Blood. 2003;102(10):3665–7. doi: 10.1182/blood-2003-02-0460.

37. Liu Y, Wada R, Yamashita T, Mi Y, Deng CX, Hobson JP, Rosenfeldt HM, Nava VE, Chae SS, Lee MJ, Liu CH, Hla T, Spiegel S, Proia RL, Edg-1, the G protein-coupled receptor for sphingosine-1-phosphate, is essential for vascular maturation, J Clin Invest, 2000; 106(8) 951–61.

38. Kalinichenko VV, Zhou Y, Shin B, Stolz DB, Watkins SC, Whitsett JA, Costa RH. Wild-type levels of the mouse Forkhead Box f1 gene are essential for lung repair. Am J Physiol Lung Cell Mol Physiol. 2002;282(6):L1253–65. doi: 10.1152/ajplung.00463.2001.

39. Kalinichenko VV, Lim L, Stolz DB, Shin B, Rausa FM, Clark J, Whitsett JA, Watkins SC, Costa RH. Defects in pulmonary vasculature and perinatal lung hemorrhage in mice heterozygous null for the Forkhead Box f1 transcription factor. Dev Biol. 2001; 235:489–506. doi: 10.1006/dbio.2001.0322.

40. Cai Y, Bolte C, Le T, Goda C, Xu Y, Kalin TV, Kalinichenko VV. FOXF1 maintains endothelial barrier function and prevents edema after lung injury. Sci Signal. 2016;9(424):ra40. doi: 10.1126/scisignal.aad1899.

41. Cartier A, Leigh T, Liu CH, Hla T. Endothelial sphingosine 1-phosphate receptors promote vascular normalization and antitumor therapy. Proc Natl Acad Sci U S A. 2020;117(6):3157–3166. doi: 10.1073/pnas.1906246117.

42. Schneider DJ, Lindsay JC, Zhou Y, Molina JG, Blackburn MR. Adenosine and osteopontin contribute to the development of chronic obstructive pulmonary disease. FASEB J. 2010;24(1):70–80. doi: 10.1096/fj.09-140772.

43. Knoops L, Renauld JC. IL-9 and its receptor: from signal transduction to tumorigenesis. Growth Factors. 2004;22(4):207–15. doi: 10.1080/08977190410001720879.

44. Garnier O, Vilgrain I. Dialogue between VE-Cadherin and Sphingosine 1 Phosphate Receptor1 (S1PR1) for Protecting Endothelial Functions. Int J Mol Sci. 2023;24(4):4018-.doi: 10.3390/ijms24044018.

45. Imenez Silva PH, Camara NO, Wagner CA. Role of proton-activated G protein-coupled receptors in pathophysiology. Am J Physiol Cell Physiol. 2022;323(2):C400–C414. doi: 10.1152/ajpcell.00114.2022.

46. Acun T, Demir K, Oztas E, Arango D, Yakicier MC. PTPRD is homozygously deleted and epigenetically downregulated in human hepatocellular carcinomas. OMICS. 2015;19(4):220–9. doi: 10.1089/omi.2015.0010.

47. Boll M, Foltz M, Rubio-Aliaga I, Daniel H. A cluster of proton/amino acid transporter genes in the human and mouse genomes. Genomics. 2003;82(1):47–56. doi: 10.1016/s0888-7543(03)00099-5.

48. Wang Z, Yemanyi F, Blomfield AK, Bora K, Huang S, Liu CH, Britton WR, Cho SS, Tomita Y, Fu Z, Ma JX, Li WH, Chen J. Amino acid transporter SLC38A5 regulates developmental and pathological retinal angiogenesis. Elife. 2022;11:e73105. doi: 10.7554/eLife.73105.

49. Amatschek S, Kriehuber E, Bauer W, Reininger B, Meraner P, Wolpl A, Schweifer N, Haslinger C, Stingl G, Maurer D. Blood and lymphatic endothelial cell-specific differentiation programs are stringently controlled by the tissue environment. Blood. 2007;109(11):4777–85. doi: 10.1182/blood-2006-10-053280.

50. Dreffs A, Lin CM, Liu X, Shanmugam S, Abcouwer SF, Kern TS, Antonetti DA. All-trans-Retinaldehyde Contributes to Retinal Vascular Permeability in Ischemia Reperfusion. Invest Ophthalmol Vis Sci. 2020;61(6):8. doi: 10.1167/iovs.61.6.8.

51. Wang M. DPEP1 mediates neutrophil and monocyte influx. Nat Rev Nephrol. 2022;18(4):199. doi: 10.1038/s41581-022-00554-3.

52. Lin YC, Sahoo BK, Gau SS, Yang RB. The biology of SCUBE. J Biomed Sci. 2023;30(1):33. doi: 10.1186/s12929-023-00925-3.

53. Carmeliet P, Lampugnani MG, Moons L, Breviario F, Compernolle V, Bono F, Balconi G, Spagnuolo R, Oosthuyse B, Dewerchin M, Zanetti A, Angellilo A, Mattot V, Nuyens D, Lutgens E, Clotman F, de Ruiter MC, Gittenberger-de Groot A, Poelmann R, Lupu F, Herbert JM, Collen D, Dejana E. Targeted deficiency or cytosolic truncation of the VE-cadherin gene in mice impairs VEGF-mediated endothelial survival and angiogenesis. Cell. 1999;98(2):147–57. doi: 10.1016/s0092-8674(00)81010-7.

54. Boggon TJ, Eck MJ. Structure and regulation of Src family kinases. Oncogene. 2004;23(48):7918–27. doi: 10.1038/sj.onc.1208081.

55. Daday C, de Buhr S, Mercadante D, Gräter F. Mechanical force can enhance c-Src kinase activity by impairing autoinhibition. Biophys J. 2022;121(5):684–691. doi: 10.1016/j.bpj.2022.01.028.

56. Gaengel K, Niaudet C, Hagikura K, Laviña B, Muhl L, Hofmann JJ, Ebarasi L, Nyström S, Rymo S, Chen LL, Pang MF, Jin Y, Raschperger E, Roswall P, Schulte D, Benedito R, Larsson J, Hellström M, Fuxe J, Uhlén P, Adams R, Jakobsson L, Majumdar A, Vestweber D, Uv A, Betsholtz C. The sphingosine-1-phosphate receptor S1PR1 restricts sprouting angiogenesis by regulating the interplay between VE-cadherin and VEGFR2. Dev Cell. 2012;23(3):587–599. doi: 10.1016/j.devcel.2012.08.005.

57. Lee MJ, Thangada S, Claffey KP, Ancellin N, Liu CH, Kluk M, Volpi M, Sha’afi RI, Hla T. Vascular endothelial cell adherens junction assembly and morphogenesis induced by sphingosine-1-phosphate. Cell. 1999;99(3):301–12. doi: 10.1016/s0092-8674(00)81661-

58. Lieu C, Heymach J, Overman M, Tran H, Kopetz S. Beyond VEGF: inhibition of the fibroblast growth factor pathway and antiangiogenesis. Clin Cancer Res. 2011;17(19):6130–9. doi: 10.1158/1078-0432.

59. West JB. Pulmonary capillary stress failure. J Appl Physiol 1985;89(6):2483–9;doi: 10.1152/jappl.2000.89.6.2483.

60. Kee K, Stuart-Andrews C, Nilsen K, Wrobel JP, Thompson BR, Naughton MT. Ventilation heterogeneity is increased in patients with chronic heart failure. Physiol Rep. 2015;3(10):e12590. doi: 10.14814/phy2.12590.

61. Hisata S, Racanelli AC, Kermani P, Schreiner R, Houghton S, Palikuqi B, Kunar B, Zhou A, McConn K, Capili A, Redmond D, Nolan DJ, Ginsberg M, Ding BS, Martinez FJ, Scandura JM, Cloonan SM, Rafii S, Choi AMK. Reversal of emphysema by restoration of pulmonary endothelial cells. J Exp Med. 2021;218(8):e20200938. doi: 10.1084/jem.20200938.

62. Hägg P, Väisänen T, Tuomisto A, Rehn M, Tu H, Huhtala P, Eskelinen S, Pihlajaniemi T. Type XIII collagen: a novel cell adhesion component present in a range of cell-matrix adhesions and in the intercalated discs between cardiac muscle cells. Matrix Biol. 2001;19(8):727–42. doi: 10.1016/s0945-053x(00)00119-0.

63. Sund M, Ylönen R, Tuomisto A, Sormunen R, Tahkola J, Kvist AP, Kontusaari S, Autio-Harmainen H, Pihlajaniemi T. Abnormal adherence junctions in the heart and reduced angiogenesis in transgenic mice overexpressing mutant type XIII collagen. EMBO J. 2001;20(18):5153–64. doi: 10.1093/emboj/20.18.5153.

64. Sakurai Y, Ohgimoto K, Kataoka Y, Yoshida N, Shibuya M. Essential role of Flk-1 (VEGF receptor 2) tyrosine residue 1173 in vasculogenesis in mice. Proc Natl Acad Sci U S A. 2005;102(4):1076–81. doi: 10.1073/pnas.0404984102.

65. Zack GW, Rogers WE, Latt SA. Automatic measurement of sister chromatid exchange frequency. J Histochem Cytochem. 1977;25(7):741–753. doi: 10.1177/25.7.70454.

66. Z. Püspöki, M. Storath, D. Sage, M. Unser Transforms and Operators for Directional Bioimage Analysis: A Survey Advances in Anatomy, Embryology and Cell Biology, 2016.vol. 219, Springer International Publishing, ch. 3.

67. Thurlbeck WM. Measurement of pulmonary emphysema. Am Rev Respir Dis. 1967;95(5):752–64. doi: 10.1164/arrd.1967.95.5.752. PMID: 5337140.

